# Age-related blunting of serial sarcomerogenesis and mechanical adaptations following 4 weeks of maximal eccentric resistance training

**DOI:** 10.1101/2023.11.07.566004

**Authors:** Avery Hinks, Makenna A. Patterson, Binta S. Njai, Geoffrey A. Power

## Abstract

**Background:** During natural aging, muscles atrophy, which is partly accounted for by a loss of sarcomeres in series. Serial sarcomere number (SSN) is associated with aspects of muscle mechanical function including the force-length and force-velocity-power relationships; hence, the age-related loss of SSN contributes to declining performance. Training emphasizing muscle lengthening (eccentric) contractions increases SSN in young healthy rodents. However, the ability for eccentric training to increase SSN and improve mechanical function in old age is unknown.

**Methods:** Ten young (9 months) and 11 old (33 months) Fisher344/BN F1 rats completed 4 weeks of unilateral isokinetic eccentric plantar flexion training 3 days/week. Pre- and post-training, the plantar flexors were assessed for maximum tetanic torque (ankle angles of 70° and 90°), the torque-frequency relationship (stimulation frequencies of 1-100 Hz), the passive torque-angle relationship (ankle angles of 110-70°), and the torque-angular velocity-power relationship (isotonic loads of 10%-80% maximum). Following post-training testing, rats were sacrificed, and the soleus, lateral gastrocnemius (LG), and medial gastrocnemius (MG) were harvested for SSN assessment by measuring sarcomere lengths with laser diffraction, with the untrained leg used as a control.

**Results:** In the untrained leg/pre-training, old rats had lower SSN in the soleus (–9%), LG (–7%), and MG (–14%), lower maximum torque (–27 to –42%), power (–63%), and shortening velocity (–35%), and greater passive torque (+62 to +191%) than young. Young rats showed increased SSN from the untrained to the trained soleus and MG. In contrast, old rats had no change in soleus SSN between legs and experienced SSN loss in the LG. Pre- to post-training, young rats saw modest improvements in isometric mechanical function, including a 13% increase in maximum torque at 90° and 4-11% increases in 10-60 Hz torque. Old rats, however, had reductions in maximum torque (–35%), shortening velocity (–46%), and power (–63%), and increased passive torque (+24 to +51%) from pre- to post-training.

**Conclusions:** Eccentric training induced serial sarcomerogenesis and improved mechanical function in young rats, while old rats exhibited dysfunctional remodeling that led to impairments in muscle mechanical performance following training.

## Background

During natural aging, skeletal muscle experiences a progressive loss of muscle mass and cross-sectional area (1–5), as well as reductions in muscle fascicle pennation angle and fascicle length (FL) (6–8). The reduced muscle FL with age has been shown in rodents to be driven by a loss of sarcomeres aligned in series (9,10). Concomitantly, aging is associated with declines in neuromuscular performance, including reduced strength and power output, a slower maximum shortening velocity, and elevated resting passive tension (5,11–13). A muscle’s serial sarcomere number (SSN) is directly proportional to its maximum excursion and maximum shortening velocity (13,14). Having more sarcomeres in series is also conducive to greater mechanical work and power during shortening contractions for a given joint excursion (14–16). As well, passive force increases exponentially with muscle stretch due in part to the viscoelastic protein titin (17), and it follows that a greater SSN can alleviate passive tension at longer muscle lengths (16,18) owing to shorter sarcomere lengths (SL). Due to these close associations between SSN and biomechanical properties of muscle, interventions promoting the addition of sarcomeres in series have been suggested as a method to mitigate age-related impairments in muscle mechanical function (7).

Training biased to active lengthening (i.e., eccentric) muscle contractions is commonly used to induce serial sarcomerogenesis in healthy young animals (18–21) and increase FL (as detected by ultrasound) in healthy young humans (22–25). There is, however, often a blunted muscle hypertrophic response to resistance training in old compared to young individuals (11,26–29), and this may extend to a dysregulation of serial sarcomere addition in old compared to young (8). Quinlan et al. (30) observed a 4% increase in vastus lateralis FL 2 weeks into moderate-load eccentric training in young men, while a comparable magnitude of FL increase did not occur until 4 weeks in older (∼68 years) men, and Raj et al. (31) observed no changes in vastus lateralis or medial gastrocnemius FL in older (∼68 years) men following 16 weeks of eccentric training. There may also be an uncoupling between adaptations in FL and adaptations in mechanical function. For example, while Thom et al. (12) showed that a 19% difference in FL between young and older (∼75 years) men accounted for almost half the age-related difference in maximum shortening velocity, Reeves et al. (32) observed no changes to the force-velocity relationship following eccentric training in older (∼67 years) men despite a 20% increase in FL. It is important to note that ultrasound was used to measure FL in these studies on humans. Since ultrasound is limited to only capturing FL and not SL, it is unclear whether any observed increases in FL (or lack thereof) were associated with an increase in SSN (33,34). Altogether, there may be an age-related impairment in eccentric training-induced serial sarcomerogenesis, but current studies using non-invasive methods in humans make this difficult to resolve.

Studies of eccentric exercise-induced muscle damage have also shown a clear deficit in the ability to recover strength and adapt following a single bout of eccentric exercise in old compared to young rodents (35,36) and humans (37–39). This inability to adapt to eccentric exercise likely arises from a combination of factors. Age-related changes in pathways controlling mechanotransduction, gene expression, and protein synthesis limit the sarcomerogenic response to mechanical stimuli (8). The stiffer extracellular matrix (ECM) in older individuals (40,41) may also limit the mechanical stimuli imposed on muscle contractile tissue, as pre-training collagen content and packing density are negatively associated with hypertrophic outcomes (42,43). As well, if aged muscle has a lower SSN to begin with (10), they may incur greater stretch and thus damage than young muscle for a given magnitude of joint rotation, leading to a longer recovery period (36,44,45). There has, however, been no direct investigation into whether older individuals can add sarcomeres in series to the same extent as young following long-term eccentric training.

The present study investigated serial sarcomerogenesis (i.e., adding sarcomeres in series) and corresponding adaptations in muscle mechanical function of the plantar flexors in old (33 months) compared to young (9 months) Fisher 344/Brown Norway F1 rats following 4 weeks of isokinetic eccentric training. We hypothesized that young rats would experience an increase in SSN, which would correspond to improvements in torque production, maximum shortening velocity, and peak power, and a reduction in resting passive tension. We hypothesized that old rats would experience a smaller magnitude of serial sarcomerogenesis compared to young rats, which would correspond to smaller alterations in mechanical function.

## Methods

### Animals

Eleven old (sacrificial age ∼33 months) and 10 young (sacrificial age ∼9 months) Fisher 344/Brown Norway F1 rats were obtained (Charles River Laboratories, Senneville, QC, Canada) with approval from the University of Guelph’s Animal Care Committee (AUP #4905) and all protocols following CCAC guidelines. Rats were housed at 23°C in groups of two or three and given ad-libitum access to a Teklad global 18% protein rodent diet (Envigo, Huntington, Cambs., UK) and room-temperature water. Pre-training mechanical testing measurements were obtained when the rats were ∼32 and ∼8 months of age for the old and young groups, respectively, then training commenced no more than 1 week later. Post-training mechanical testing was completed 48-72 hours following the final training session, then the hindlimbs were fixed in formalin for subsequent SSN estimations from the soleus, lateral gastrocnemius (LG), and medial gastrocnemius (MG). The left leg completed training while the right leg acted as an internal control for SSN (34,46–48).

### Data acquisition during mechanical testing and training

A 701C High-Powered, Bi-Phase Stimulator (Aurora Scientific, Aurora, ON, Canada) was used to evoke transcutaneous muscle stimulation via a custom-made electrode holder with steel galvanized pins situated transversely over the popliteal fossa and the calcaneal tendon (Figure 1A). During piloting, we determined this stimulation setup produced similar values of maximum isometric tetanic torque across repeated testing sessions. Torque, angle, and stimulus trigger data were sampled at 1000 Hz with a 605A Dynamic Muscle Data Acquisition and Analysis System (Aurora Scientific, Aurora, ON, Canada) with a live torque tracing during all training and mechanical data collection sessions.

**Figure 1:**
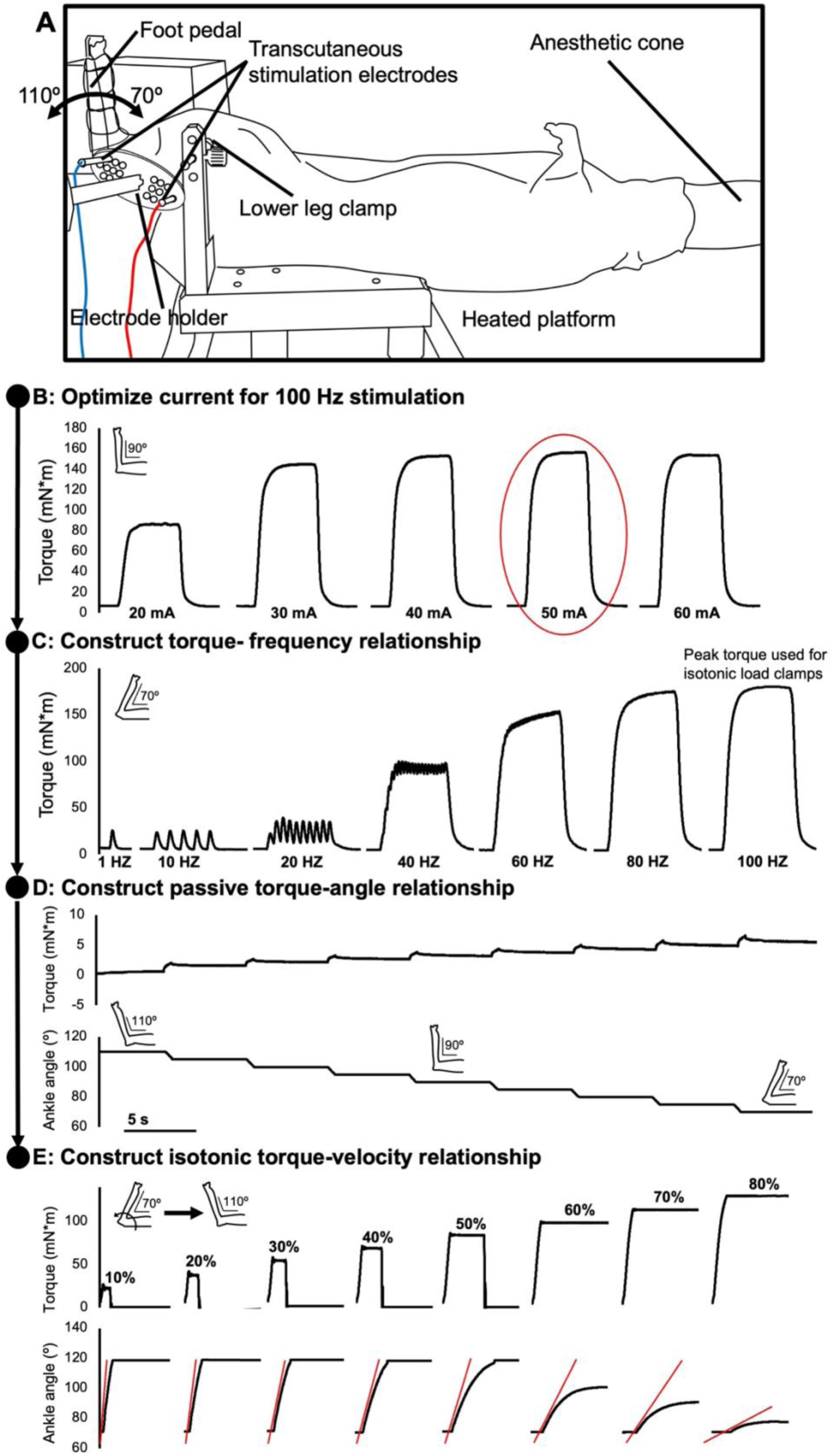
Experimental setup and timeline. A: The rat’s left foot was secured to a foot pedal and electrically stimulated to evoke plantar flexion contractions, with the knee fixed at 90°. The ankle had a 70° to 110° range of motion. B: With the ankle at 90°, the optimal current for 100 Hz stimulation was determined by ramping up in 10 mA increments. C: With the ankle at 70°, the torque-frequency relationship was constructed in order from 1 Hz to 100 Hz. D: The passive torque-angle relationship was then constructed throughout at 10° increments throughout the range of motion. E: 100 Hz isotonic contractions from 70° to 110° were performed at load clamps of 10% to 80% of maximum isometric torque at 70°, in a randomized order. Red lines represent the maximum derivative of the position-time trace (i.e., the measurement of isotonic shortening velocity).

### Mechanical testing

The rats were anesthetized with isoflurane and positioned on a heated platform (37°C) in a supine position. After shaving the leg completely of hair, the left leg was fixed to a force transducer/length controller foot pedal via tape, with the knee immobilized at 90° (Figure 1A). Each mechanical testing session began with determination of the optimal current for stimulation (frequency = 100 Hz, pulse duration = 0.1 ms, train duration = 500 ms) at an ankle angle of 90° (full plantar flexion = 180°), which was the current used throughout the remainder of the session (Figure 1B). The dorsiflexor muscles were observed visually and palpated to confirm there was minimal antagonist activation during tetanic contractions. A torque-frequency relationship was then constructed with 500-ms isometric contractions at stimulation frequencies of 1, 10, 20, 40, 60, 80, and 100 Hz (Figure 1C). These were completed at an ankle angle of 70° because during piloting, we found that this ankle angle produced the highest active plantar flexion torque among the angles available with the Aurora system (70° to 110°). The 100-Hz contraction was used for determination of relative load clamps for the subsequent isotonic contractions. Active torque was measured by subtracting the minimum value of torque at baseline (i.e., the passive torque) from the maximum value of total torque during stimulation (18,49). A passive torque-angle relationship was then constructed by recording the minimum passive torque following 5 s of stress-relaxation at ankle angles of 110, 105, 100, 95, 90, 85, 80, 75, and 70° (Figure 1D). A torque-angular velocity-power relationship was then constructed from isotonic contractions. For each isotonic contraction, the ankle started at 70°, then contracted against the given load clamp to a maximum angle of 110° (Figure 1E). Isotonic contractions were performed at load clamps equating to 10, 20, 30, 40, 50, 60, 70, and 80% of the maximum 100-Hz isometric torque at 70°, in a randomized order. Angular shortening velocity was recorded as the maximum time derivative of the angular displacement during the isotonic contraction. Two minutes of rest were provided between each stimulation to minimize the development of muscle fatigue.

To determine the frequency at which half of maximum active torque was achieved (F_50_), torque-frequency data were fitted the following equation (50):

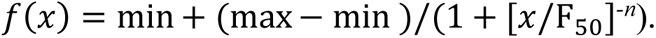

Where *x* is the stimulation frequency, min and max are the respective minimum and maximum torque values recorded across the torque-frequency curve, and *n* is the coefficient describing the slope of the steepest portion of the torque-frequency curve.

The estimated maximum shortening velocity at zero load (V_max_), maximum power (i.e., torque multiplied by angular velocity), the torque and velocity values that corresponded to peak power, and the curvature of the torque-velocity relationship (*a* from the equation below divided by the maximum 100-Hz isometric torque at 70°) were determined by fitting the measured torque and angular velocity values to a rectangular hyperbolic curve using the equation (51):

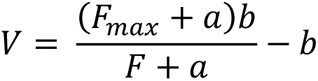

With F_max_ being the maximum 100-Hz isometric torque at 70°, *F* and *V* representing the isotonic load clamp’s torque and the angular velocity, respectively, and *a* and *b* representing Hill’s thermodynamic constants with the units of torque (N•m) and velocity (°/s), respectively.

All curve-fitting was performed in Python 3 using a least square error optimization code.

### Isokinetic eccentric training

The present study’s eccentric training protocol was adapted from previous investigations in rats (52,53) and rabbits (21) that have been shown to induce muscle morphological adaptations, including (for the rabbit study) increases in SSN. Training lasted 4 weeks and occurred 3 days/week (Monday, Wednesday, Friday) (18,21). For the training, rats were anesthetized with isoflurane and positioned on a heated platform (37°C) in a supine position. The left foot was fixed to a force transducer/length controller with the knee immobilized at 90°. At the start of each session, the stimulus current was first adjusted to produce maximum tetanic torque (pulse duration = 0.1 ms, frequency = 100 Hz, train duration = 500 ms) (21,53,54) at an ankle angle of 90° (i.e., the ankle angle at the centre of the range of motion during the eccentric contractions).

Each eccentric repetition, during maximal activation, consisted of 3 phases: 1) a 500-ms pre-activation at an ankle angle of 110° (the maximum angle of the Aurora system); 2) active lengthening to 70° (the minimum angle of the Aurora system); and 3) 3 seconds of deactivation followed by a return to 110° at 20 °/s. An additional 3 seconds of rest were provided between repetitions (i.e., 8 seconds total between each stimulation). This 70° to 110° range of motion corresponds closely to that of the ankle during the stance phase of rodent voluntary ambulation (55). To progress the eccentric training stimulus throughout the 4 weeks, we increased the number of repetitions and the velocity of the eccentric contractions. Increasing velocity in particular is typically associated with greater eccentric force development and a stronger stimulus for serial sarcomere addition (56). During week 1, rats completed 3 sets of 8 repetitions at 40°/s. In week 2, they completed 3 sets of 9 repetitions at 40°/s. Weeks 3 and 4 consisted of 3 sets of 9 and 10 repetitions, respectively, at 80°/s. Two minutes of rest were provided between each set. Maximum isometric torque during the stimulation current optimization was recorded for each training session.

### Serial sarcomere number estimations

Following the post-training mechanical testing, rats were sacrificed via isoflurane anesthetization followed by CO_2_ asphyxiation. The hindlimbs were amputated and fixed in 10% phosphate-buffered formalin with the ankle pinned at 90° and the knee fully extended. After fixation for 1-2 weeks, the muscles were dissected and rinsed with phosphate-buffered saline. The muscles were then digested in 30% nitric acid for 6-8 hours to remove connective tissue and allow for individual muscle fascicles to be teased out (18,20).

For each muscle, two fascicles were obtained from each of the proximal, middle, and distal regions of the muscle (i.e., n = 6 fascicles total per muscle) and averaged for each region for the reporting of data (i.e., n = 1 averaged measurement at each of the proximal, middle, and distal regions). Dissected fascicles were placed on glass microslides (VWR International, USA), then FLs were measured using ImageJ software (version 1.53f, National Institutes of Health, USA) from pictures captured by a level, tripod-mounted digital camera, with measurements calibrated to a ruler in plane with the fascicles. Sarcomere length measurements were taken at n = 6 different locations proximal to distal along each fascicle via laser diffraction (Coherent, Santa Clara, CA, USA) with a 5-mW diode laser (25 μm beam diameter, 635 nm wavelength) and custom LabVIEW program (Version 2011, National Instruments, Austin, TX, USA) (57), for a total of n = 36 sarcomere length measurements per muscle. For each fascicle, the six SL measurements were averaged to obtain a value of average SL. Serial sarcomere numbers was calculated as:

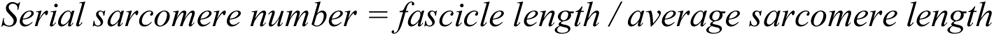

Within each fascicle, SL standard deviation (SL SD) was also noted as an estimate of SL non-uniformity.

### Statistical analysis

All statistical analyses were performed in SPSS Statistics Premium 28. To compare the torque-frequency relationship between young and old rats and pre- to post-training, a three-way repeated measures analysis of variance (ANOVA) was used with training (pre-training, post-training) and frequency (1, 10, 20, 40, 60, 80, 100 Hz) as within-subjects factors and age (old, young) as a between-subjects factor. To compare the passive torque-angle relationship between young and old rats and pre- to post-training, a three-way repeated measures ANOVA was used with training (pre-training, post-training) and angle (110°, 105°, 100°, 95°, 90°, 85°, 80°, 75°, 70°) as within-subjects factors and age (old, young) as a between-subjects factor. While a full active torque-angle relationship was not constructed, we obtained active torque at two ankle angles: 90° (during optimization of the stimulation current) and 70° (when constructing the torque-frequency relationship). Therefore, to gain insight into the active torque-angle relationship compared between young and old rats and pre- to post-training, a three-way repeated measures ANOVA was used with training (pre-training, post-training) and angle (90°, 70°) as within-subjects factors and age (old, young) as a between-subjects factor. To compare velocity as a function of load between young and old rats and pre- to post-training, a three-way repeated measures ANOVA was used with training (pre-training, post-training) and load (10, 20, 30, 40, 50, 60, 70, 80%) as within-subjects factors and age (old, young) as a between-subjects factor. To investigate changes in maximum torque at 90° throughout the training period, we used a three-way repeated measures ANOVA with training (pre-training, post-training) and day (pre, day 1, day 2, day 3, day 4, day 5, day 6, day 7, day 8, day 9, day 10, day, 11, day 12, post) as within-subjects factors and age (old, young) as a between-subjects factor. Two-way repeated measures ANOVAs with training (pre-training, post-training) as a within-subjects factor and age (old, young) as a between-subjects factor were used to compare the following functional measures between young and old rats and pre- to post-training: body mass, F_50_, the *n* coefficient, twitch half-relaxation time, V_max_, peak power, torque at peak power, velocity at peak power, curvature of the torque-velocity relationship, and the *a* and *b* coefficients of the torque-velocity relationship.

Two-way repeated measures ANOVAs with training (trained leg, untrained leg) as a within-subjects factor and age (old, young) as a between-subjects factor were used to compare muscle wet weight between legs and between young and old rats. Three-way repeated measures ANOVAs with training (trained leg, untrained leg) and muscle region (proximal, middle, distal) as within-subjects factors and age (old, young) as a between-subjects factor were used to compare SSN, FL, SL, and SL SD between legs, across muscle region, and between young and old rats.

A Greenhouse-Geisser correction was applied to all ANOVAs. Where main effects or interactions were detected, two-tailed t-tests were used for pairwise comparisons, with a Sidak correction for multiplicity. Significance was set at *P* < 0.05. Where significance was observed for main effects and interactions, the effect size was reported as the partial eta squared (η_p_^2^).

## Results

### Differences in body mass between young and old rats and pre- to post-training

There was a training × age interaction (F(1,19) = 56.724, *P* < 0.001, η_p_^2^ = 0.749) for body mass. Old rats weighed more than young both pre- (young: 401.76 ± 13.63 g; old: 531.53 ± 39.72 g; *P* < 0.001) and post-training (young: 381.61 ± 15.71 g; old: 480.79 ± 37.91 g; *P* < 0.001). As well, young (*P* < 0.001) and old rats (*P* < 0.001) both lost body mass from pre- to post-training, with old rats losing relatively more (−10%) than young (−5%).

### Differences in muscle wet weight between young and old and between the trained and untrained leg

Full ANOVA results for muscle wet weight are shown in Supplemental Figure S1. For the soleus and MG, there were effects of training (soleus: F(1,19) = 14.621, *P* = 0.015, η_p_^2^ = 0.273; MG: F(1,19) = 16.677, *P* < 0.001, η_p_^2^ = 0.467) and age (soleus: F(1,19) = 21.476, *P* <0.001, η_p_^2^ = 0.531; MG: F(1,19) = 109.960, *P* < 0.001, η_p_^2^ = 0.853) on muscle wet weight (Figure 2A-B and E-F). The trained soleus and MG weighed 8% and 10% more, respectively, than those of the untrained leg. The soleus and MG of old rats weighed 14% and 28% less, respectively, than those of young rats. There were no differences in LG wet weight between legs (F(1,19) = 1.367, *P* = 0.257), but the LG of old rats weighed 27% less than the LG of young rats (F(1,19) = 51.681, *P* < 0.001, η_p_^2^ = 0.731) (Figure 2C-D). There were no interactions of training × age on muscle wet weight.

**Figure 2:**
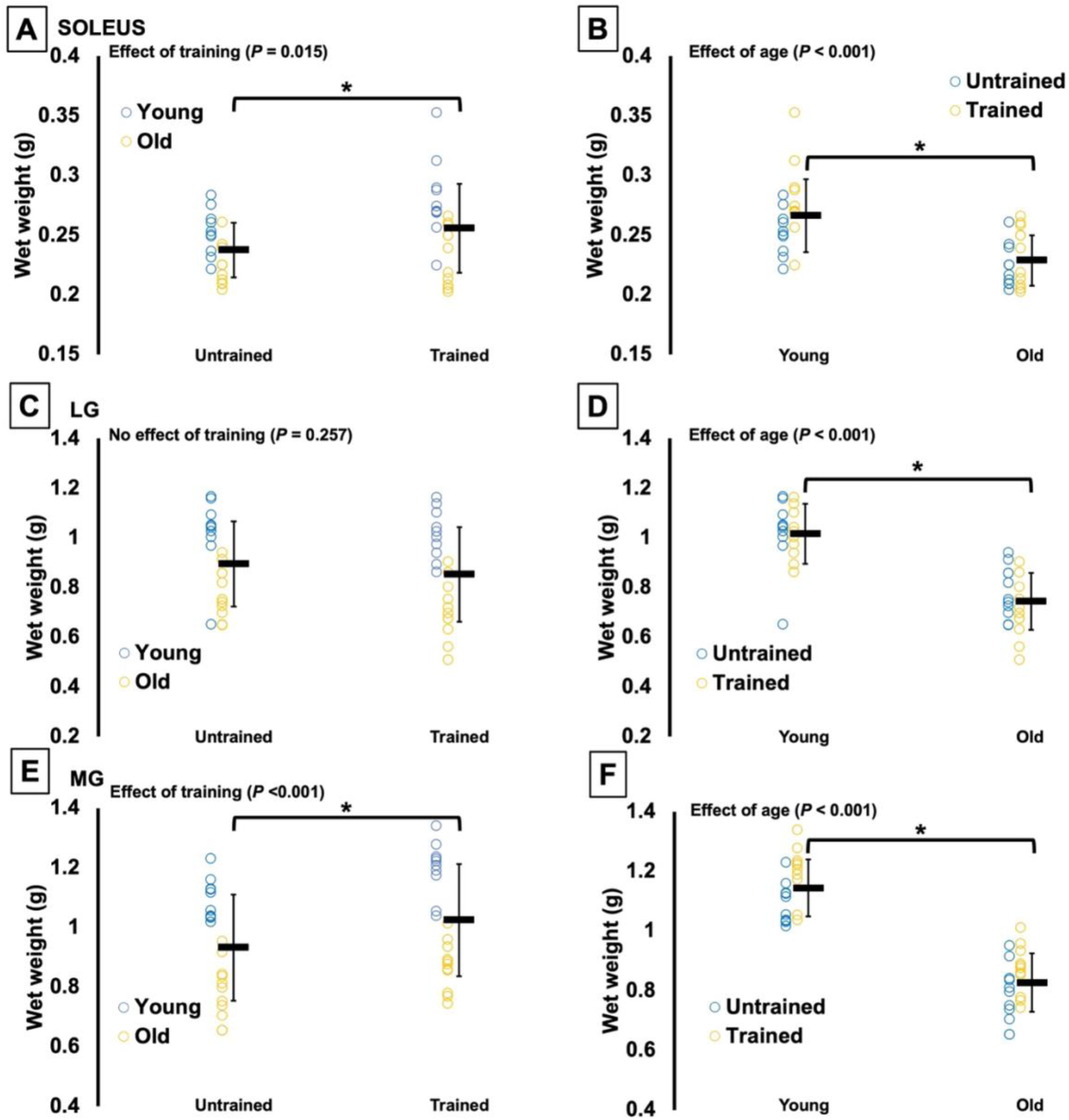
Differences in muscle wet weight of the soleus (**A and B**), lateral gastrocnemius (LG) (**C and D**) and medial gastrocnemius (**E and F**) between the trained and untrained leg (**A, C, and E**) and between young and old rats (**B, D, and F**). *Significant difference between indicated points (P < 0.05). Data are presented as mean ± standard deviation (n = 10 young and n = 11 old).

### Differences in SSN, FL, and SL between young and old, between the trained and untrained leg, and across muscle region

Full ANOVA results for SSN, FL, and SL are shown in Supplemental Figure S2.

For soleus SSN, there was a training × age interaction (F(1,19) = 10.099, *P* = 0.005, η_p_^2^ =0.347) and no effect of region (F(1.642,31.204) = 0.531, *P* = 0.592) (Figure 3A-B). In young rats, SSN of the trained soleus was 4% greater than the untrained soleus (*P* = 0.005). Old rats had no difference in SSN between the trained and untrained soleus (*P* = 0.231). Soleus SSN of old rats was less than young rats in both the untrained (–9%) and trained (–15%) legs (*P* < 0.001).

**Figure 3:**
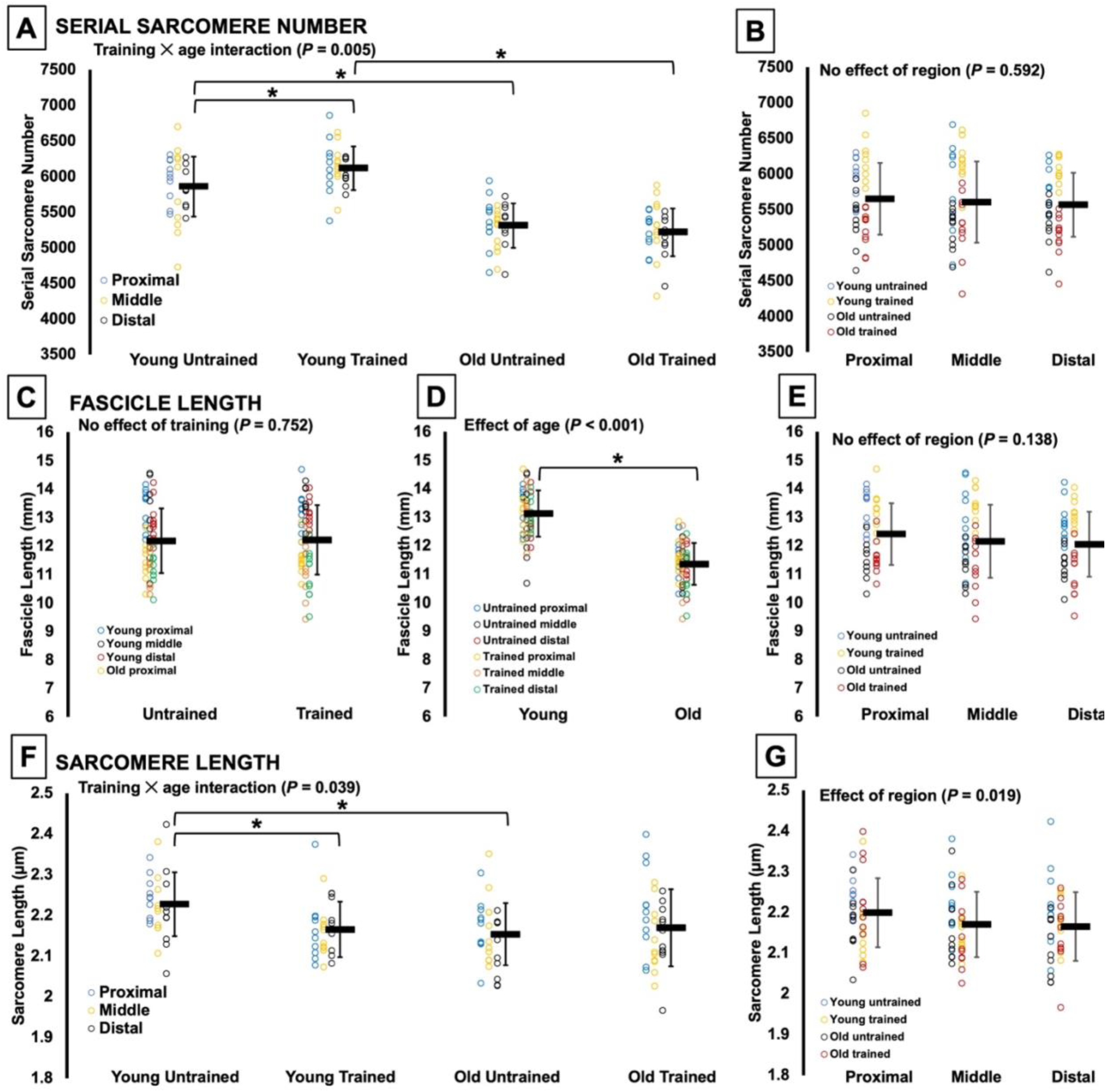
Soleus serial sarcomere number (**A and B**), fascicle length (**C-E**) and sarcomere length (**F and G**) in young and old rats in the trained and untrained legs of young and old rats across the proximal, middle, and distal regions of the muscle. *Significant difference between indicated points (P < 0.05). Data are presented as mean ± standard deviation (n = 10 young and n = 11 old).

For soleus FL, there were no effects of training (F(1,19) = 0.103, *P* = 0.752) or region (F(1.536,29.185) = 2.090, *P* = 0.138) (Figure 3C and E), but an effect of age (F(1,19) = 120.350, *P* < 0.001) showed a 13% shorter soleus FL in old than young rats (Figure 3D).

There was also a training × age interaction (F(1,19) = 4.896, *P* = 0.039, η_p_^2^ = 0.205) for soleus SL. In young rats, SL of the trained soleus was 3% shorter than the untrained soleus (*P* = 0.025), and there was no difference in soleus SL between the trained and untrained leg in old rats (*P* = 0.516). SL of the untrained soleus was 3% shorter in old compared to young rats (*P* = 0.015) (Figure 3F). There was also an effect of region (F(1.921,36.493) = 4.486, *P* = 0.019, η_p_^2^ = 0.191) on soleus SL, however, pairwise comparisons did not reveal any significant differences between regions (*P* = 0.054-0.969) so this effect seemed to be small (Figure 3G).

In summary, soleus SSN increased from the untrained to trained leg in young but not old rats. The soleus SSN increase in young rats corresponded to shorter SLs. Old rats also had shorter FLs and SLs than young rats.

For LG SSN, there was a training × age interaction (F(1,19) = 7.831, *P* = 0.011, η_p_^2^ = 0.292). In young rats, SSN did not differ between the trained and untrained LG (*P* = 0.091), but in old rats, SSN was 5% lower in the trained than untrained LG (*P* = 0.042) (Figure 4A). LG SSN of old rats was less than young in both the untrained (–7%) and trained (–16%) leg (*P* < 0.001-0.008) (Figure 4A).

**Figure 4:**
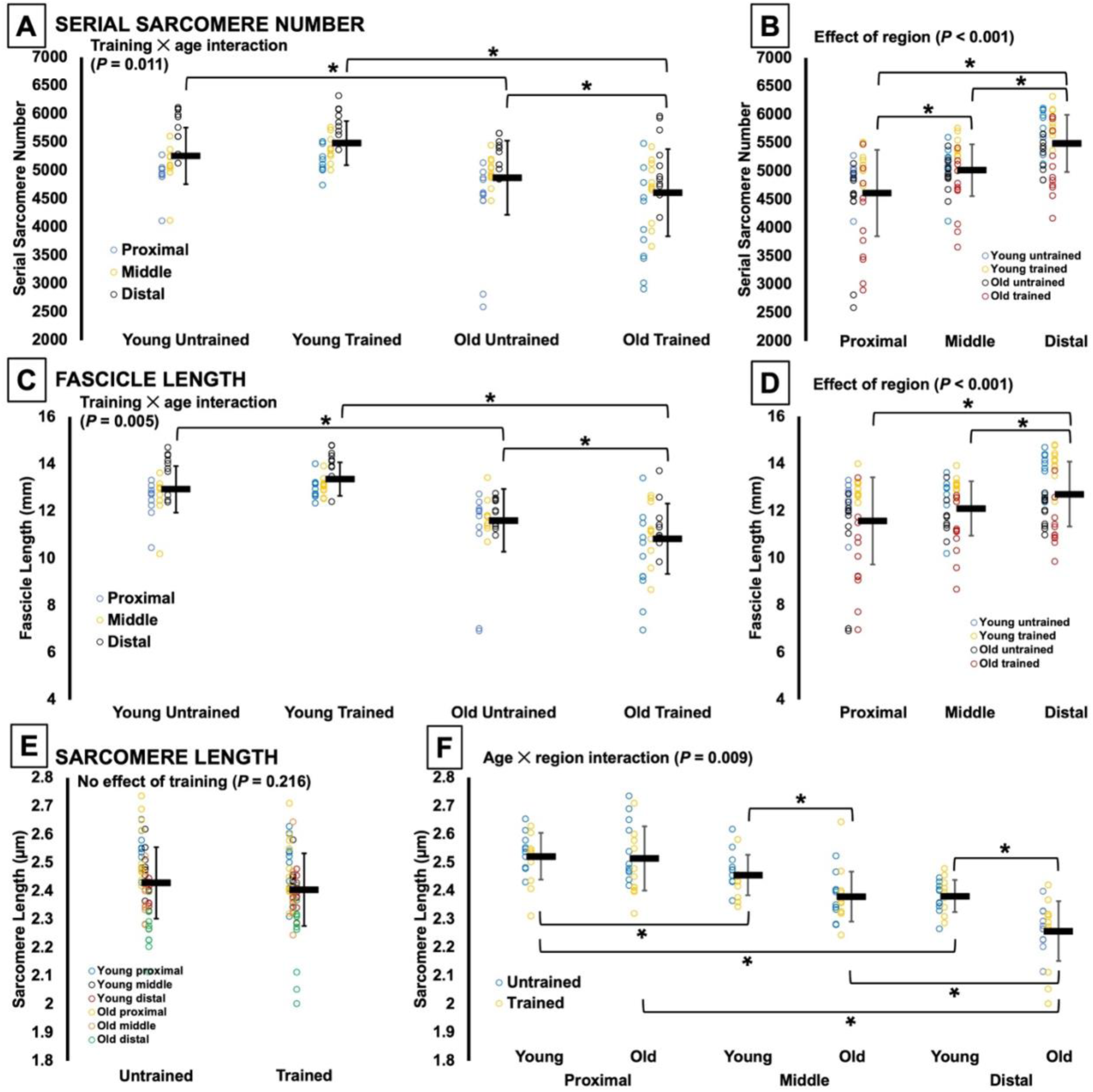
Lateral gastrocnemius serial sarcomere number (**A and B**), fascicle length (**C and D**) and sarcomere length (**E and F**) in the trained and untrained legs of young and old rats across the proximal, middle, and distal regions of the muscle. *Significant difference between indicated points (P < 0.05). Data are presented as mean ± standard deviation (n = 10 young and n = 11 old).

For LG FL, there was also a training × age interaction (F(1,19) = 10.238, *P* = 0.005, η_p_^2^ = 0.350). Like with SSN, LG FL did not differ between the trained and untrained LG for young rats (*P* = 0.130), but in old rats, FL was 7% shorter in the trained than untrained LG (*P* = 0.008) (Figure 4C). LG FL of old rats was shorter than young in both the untrained (–10%) and trained (–19%) leg (*P* < 0.001) (Figure 4C).

For LG SL, there was no effect of training (F(1,19) = 1.637, *P* = 0.216) (Figure 4E), but there was an age × region interaction (F(1.722,32.726) = 5.805, *P* = 0.009, η_p_^2^ = 0.234). Old rats had shorter LG SLs in middle (–3%) and distal (–5%) fascicles (*P* < 0.001-0.025). In young and old rats, SL decreased from proximal to distal (*P* < 0.001-0.048) (Figure 4F).

There were also effects of region on LG SSN (F(1.574,29.905) = 36.693, *P* < 0.001, η_p_^2^ = 0.659) and FL (F(1.385,26.308) = 11.495, *P* < 0.001, η_p_^2^ = 0.377) such that SSN (*P* < 0.001-0.006) and FL (*P* < 0.001-0.001) increased from proximal to distal (Figure 4B and D).

In summary, SSN and FL were not different between the untrained and trained legs for young rats, but old rats had a lower SSN and shorter FL in the trained than the untrained leg. SL did not differ between the trained and untrained legs. Old rats also had shorter SLs in the middle and distal regions of the LG compared to young rats.

For MG SSN, there was an effect of training (F(1,19) = 7.823, *P* = 0.011, η_p_^2^ = 0.292) such that SSN was 4% greater in the trained than the untrained leg (Figure 5A). There was also an effect of age (F(1,19) = 36.312, *P* < 0.001, η_p_^2^ = 0.656) such that MG SSN was 14% less in old compared to young rats (Figure 5B).

**Figure 5:**
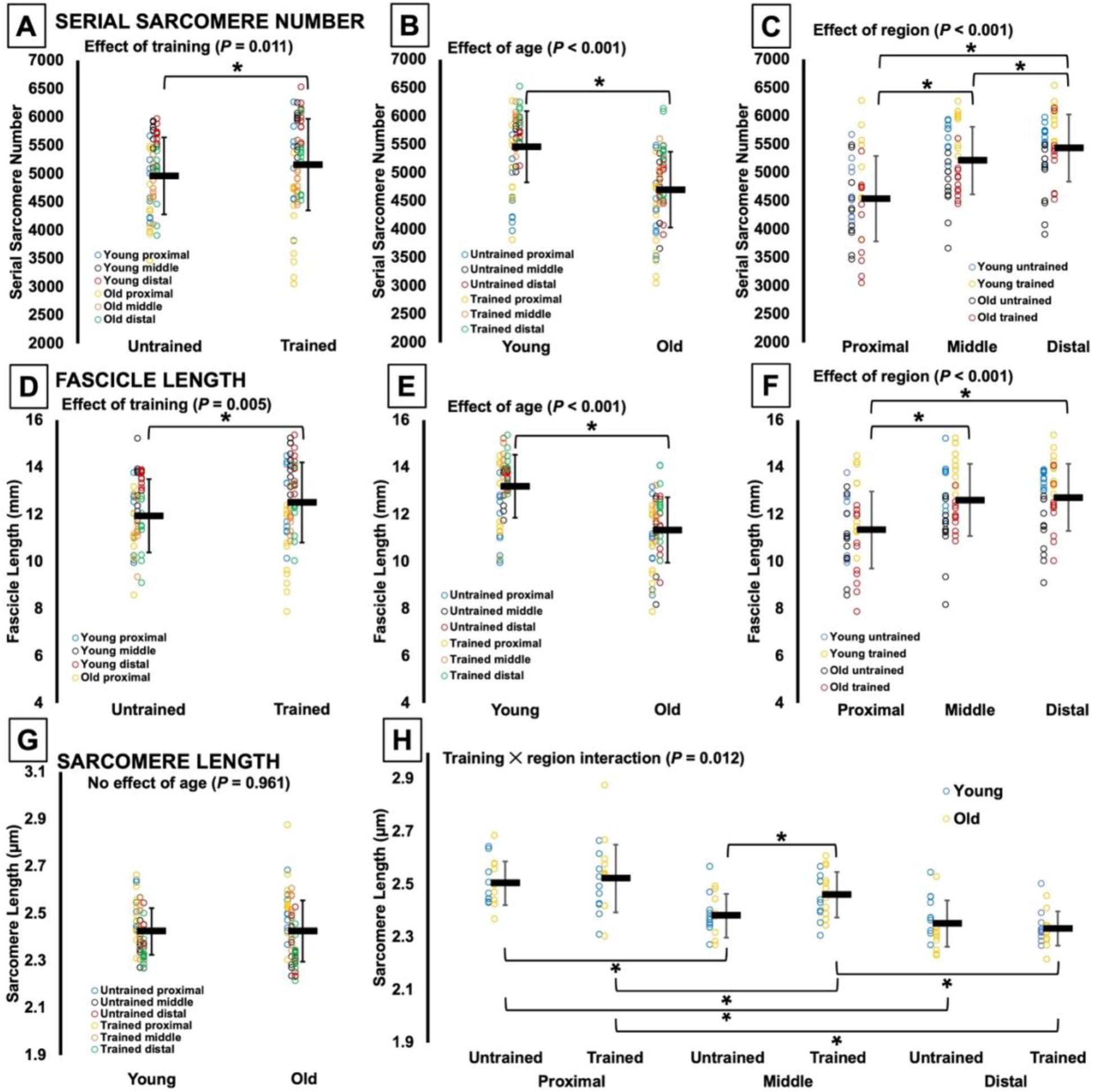
Medial gastrocnemius serial sarcomere number (**A-C**), fascicle length (**D-F**) and sarcomere length (**G and H**) in the trained and untrained legs of young and old rats across the proximal, middle, and distal regions of the muscle. *Significant difference between indicated points (P < 0.05). Data are presented as mean ± standard deviation (n = 10 young and n = 11 old).

For MG FL, there was also an effect of training (F(1,19) = 10.107, *P* = 0.005, η_p_^2^ = 0.347) such that FL was 5% longer in the trained than the untrained leg (Figure 5D). There was also an effect of age (F(1,19) = 51.640, *P* < 0.001, η_p_^2^ = 0.731) such that MG FL was 14% shorter in old compared to young rats (Figure 5E).

For MG SL, there was no effect of age (F(1,19) = 0.002, *P* = 0.961) (Figure 5G), but there was a training × region interaction (F(1.807,34.335) = 5.267, *P* = 0.012, η_p_^2^ = 0.217). MG SL was 3% longer in the trained than the untrained leg for middle fascicles (*P* = 0.006) but not proximal or distal fascicles (*P* = 0.381-0.622) (Figure 5H). SL also decreased from proximal to distal in both the trained and untrained legs (*P* <0.001-0.035) (Figure 5H).

There were also effects of region on MG SSN (F(1.413,26.843) = 31.074, *P* < 0.001, η_p_^2^ = 0.621) and FL (F(1.451,27.578) = 16.340, *P* < 0.001, η_p_^2^ = 0.462) such that SSN (*P* < 0.001-0.046) and FL (*P* < 0.001-0.002) increased from proximal to distal (Figure 5C and F).

In summary, SSN and FL of the MG increased from the untrained to the trained leg and were lower in old than young rats. SL did not differ between young and old rats but was longer in the middle region of the trained compared to the untrained leg.

### Changes in maximum strength between young and old and pre- to post-training

Full ANOVA results for 100 Hz isometric torque at 90° and 70° ankle angles are displayed in Supplemental Figure S3. There was a training × age × angle interaction (F(1,19) = 21.785, *P* < 0.001, η_p_^2^ = 0.534) for 100 Hz isometric torque. Old rats were weaker than young at both angles pre (–27% at 90° and –42% at 70°) and post-training (–60% at 90° and –65% at 70°) (*P* < 0.001) (Figure 6). Pre-training, young rats produced 8% greater torque at 70° compared to 90° (*P* < 0.001), but post-training, there was no difference in isometric torque between 70° and 90° (*P* = 0.856), with a 13% increase in torque at 90° compared to pre-training (*P* < 0.001) (Figure 6). Old rats produced 12-13% greater torque at 90° than 70° both pre (*P* < 0.001) and post-training (*P* = 0.002) and exhibited a 35% reduction in isometric torque at both angles post-training compared to pre-training (*P* < 0.001) (Figure 6).

**Figure 6:**
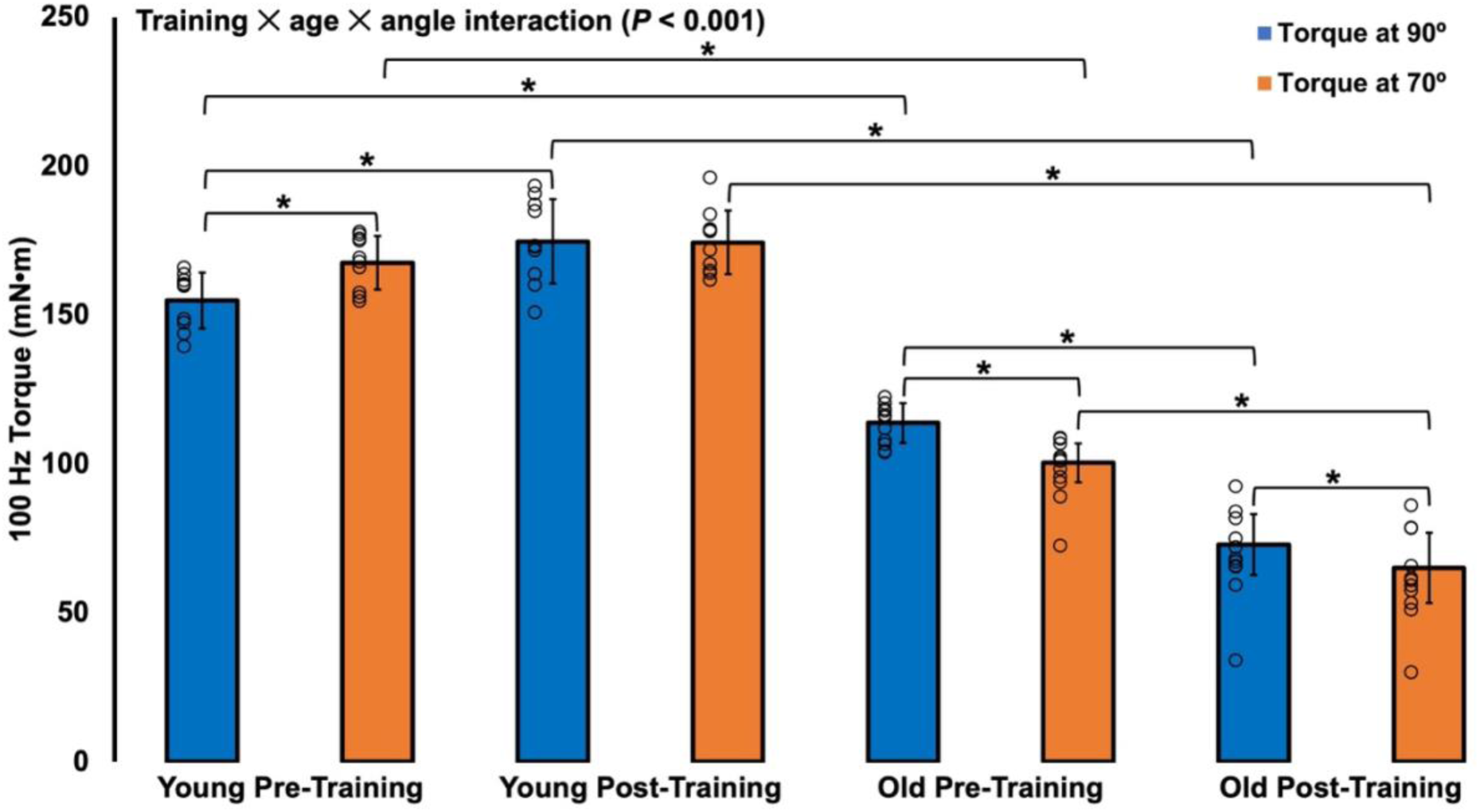
100 Hz isometric active torque at 90° and 70° ankle angles pre- and post training in young (n = 10) and old rats (n = 11). *Significant difference between indicated points (*P < 0.05*). Data are presented as mean ± standard deviation.

Full ANOVA results for specific torque (active torque/muscle mass) are displayed in Supplemental Figure S3. When normalized to the total muscle wet weight of the soleus, LG, and MG combined (post-training normalized to the trained leg, pre-training normalized to the untrained leg) to investigate how differences in muscle mass impact strength, there was a training × age × angle interaction (F(1,19) = 19.868, *P* < 0.001, η_p_^2^ = 0.511) for 100 Hz isometric torque. Pre-training, normalizing to muscle wet weight eliminated the age-related difference in torque at 90° (*P* = 0.404), and at 70° reduced the difference in torque between old and young such that old were 23% weaker (*P* < 0.001) (Figure 7). Post-training, old rats produced 45% and 51% lower specific torque than young at 90° (*P* < 0.001) and 70° (*P* < 0.001), respectively (Figure 7). Specific torque did not change from pre- to post-training for young rats at either angle (*P* = 0.165-0.690), but for old rats decreased by 40% (*P* < 0.001) (Figure 7). Therefore, differences in muscle mass accounted for some of the strength difference between young and old rats pre-training, and old rats lost strength post-training independent of changes (or lack of changes) in muscle mass.

**Figure 7:**
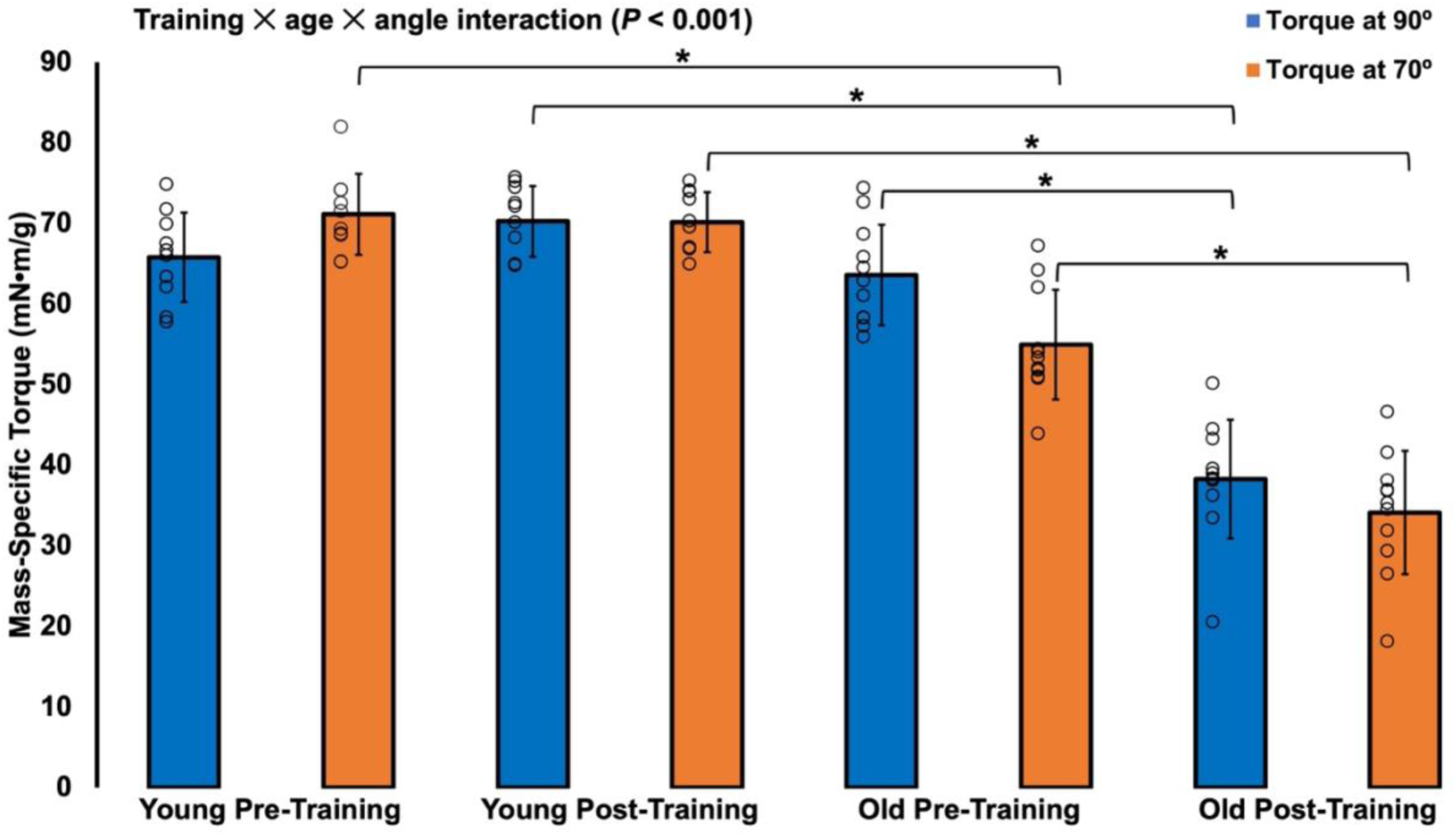
100 Hz isometric active torque at a 90° ankle angle normalized to the combined muscle wet weight of the soleus, lateral gastrocnemius, and medial gastrocnemius pre- and post-training in young (n = 10) and old rats (n = 11) (post-training normalized to the trained leg, pre-training normalized to the untrained leg). *Significant difference between indicated points (*P* < 0.05). Data are presented as mean ± standard deviation.

### Changes in the torque-frequency relationship between young and old and pre- to post-training

Full ANOVA results for the torque-frequency relationship are displayed in Supplemental Figure S4, and full ANOVA results for F_50_, the *n* coefficient, and twitch half-relaxation time are displayed in Supplemental Figure S5.

There was a training × age × frequency interaction (F(1.606,30.513) = 51.410, *P* < 0.001, η_p_^2^ = 0.730) for peak torque. Pre-training, old rats produced 14% greater torque than young at 20 Hz (*P* = 0.004), however, old rats produced 10-67% lower torque than young at all frequencies from 40 to 100 Hz (*P* = 0.014 to <0.001) (Figure 8A). Post-training, old rats produced 49-168% less torque than young at all stimulation frequencies (*P* < 0.001) (Figure 8A). Young rats produced 4-11% greater torque at frequencies from 10 to 60 Hz post-training compared to pre-training (*P* = 0.003-0.018) (Figure 8A). Old rats produced 35-38% lower torque at all frequencies post-training compared to pre-training (*P* < 0.001) (Figure 8A).

**Figure 8:**
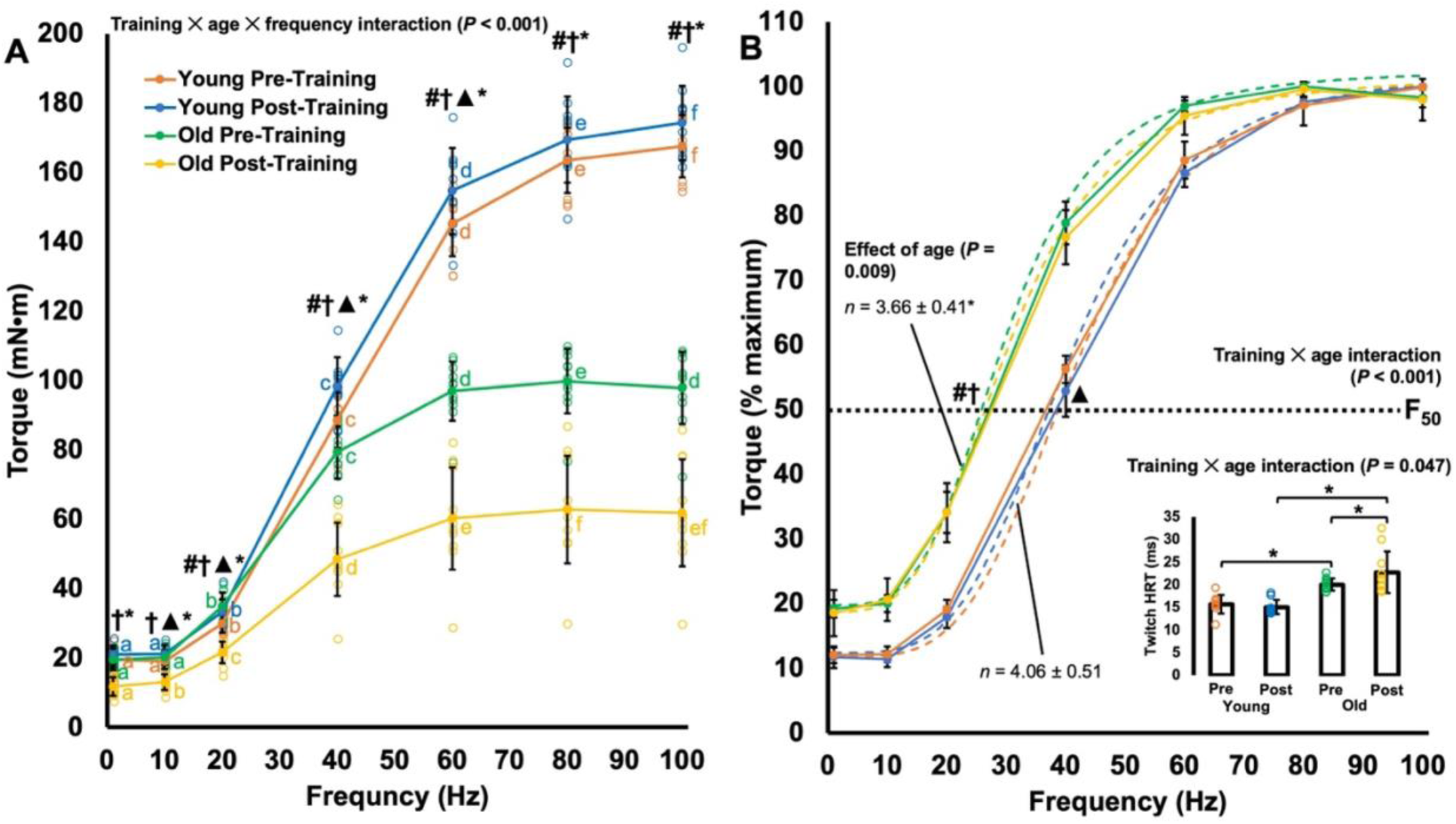
The torque-frequency relationship. **A:** Absolute experimental data with individual data points for young (n = 10) and old rats (n = 11) pre- and post-training. #Significant difference between young and old at pre-training (*P < 0.05*). †Significant difference between young and old at post-training (*P* < 0.05). ▴Significant difference between young pre- and post-training (*P* < 0.05). *Significant difference between old pre- and post-training (*P* < 0.05). Same letters indicate no significant difference between points within each color (*P* > 0.05). **B:** Torque normalized to maximum alongside the fitted curves (dashed lines). The dotted black line marks where torque is 50% of maximum (F_50_). The *n* coefficients (i.e., the Hill slope coefficient), representing the slope of the steepest part of the curve, are also displayed for old and young rats. #Significant difference in F_50_ between young and old at pre-training (*P* < 0.05). †Significant difference in F_50_ between young and old at post-training (*P* < 0.05). ▴Significant difference in F_50_ between young pre- and post-training (*P* < 0.05). *Significant effect of age for the *n* coefficient. The inset shows twitch half-relaxation time (HRT); *significant difference between indicated points (*P* < 0.05). Data are presented as mean ± standard deviation.

A training × age interaction for F_50_ (F(1,19) = 12.021, *P* = 0.003, η_p_^2^ = 0.388) also showed that F_50_ for old rats occurred at a lower frequency (pre: 29.82 ± 2.09 Hz; post: 30.42 ± 2.34 Hz) compared to young (pre: 41.84 ± 1.76 Hz; post: 40.33 ± 0.98 Hz) both pre- and post training (*P* < 0.001) (Figure 8B). Additionally, F_50_ of young rats was lower post-training (40.33 ± 0.98 Hz) compared to pre-training (41.84 ± 1.76 Hz) (*P* = 0.003) (Figure 8B). Old rats’ torque-frequency relationship also had a less steep slope (*n* coefficient) than young rats (F(1,19) = 8.436, *P* = 0.009, η_p_^2^ = 0.307) (Figure 8B).

There was also a training × age interaction for twitch half-relaxation time (F(1,19) = 4.499, *P* = 0.047, η_p_^2^ = 0.191). Old rats had a longer half-relaxation time than young both pre (+25%; *P* < 0.001) and post-training (+53%; *P* < 0.001) (Figure 8B). As well, while young rats did not differ pre- to post-training (*P* = 0.584), old rats had a 15% longer half-relaxation time post-training than pre-training (*P* = 0.022) (Figure 8B).

### Changes in maximum torque during the training period

Full ANOVA results for 100 Hz torque at 90° throughout the training period are displayed in Supplemental Figure S6. There was a day × age interaction (F(4.393,79.069) = 29.110, *P* < 0.001, η_p_^2^ = 0.618) for 100 Hz isometric torque at 90° throughout the 4-week training period. Old rats produced lower torque than young throughout the entire training period (*P* < 0.001). Young rats exhibited at first a reduction in torque throughout the first week, likely owing to eccentric exercise-induced muscle damage, differing from pre-training at day 3 (*P* = 0.008) (Figure 9), but gradually recovered thereafter, eventually improving on pre-training values post-training (Figure 6). Old rats, conversely, experienced a gradual reduction in torque throughout the training period, differing from pre-training values at days 11 and 12 in addition to post-training (*P* < 0.001-0.009) (Figure 9), likely indicating a prolonged state of muscle damage.

**Figure 9:**
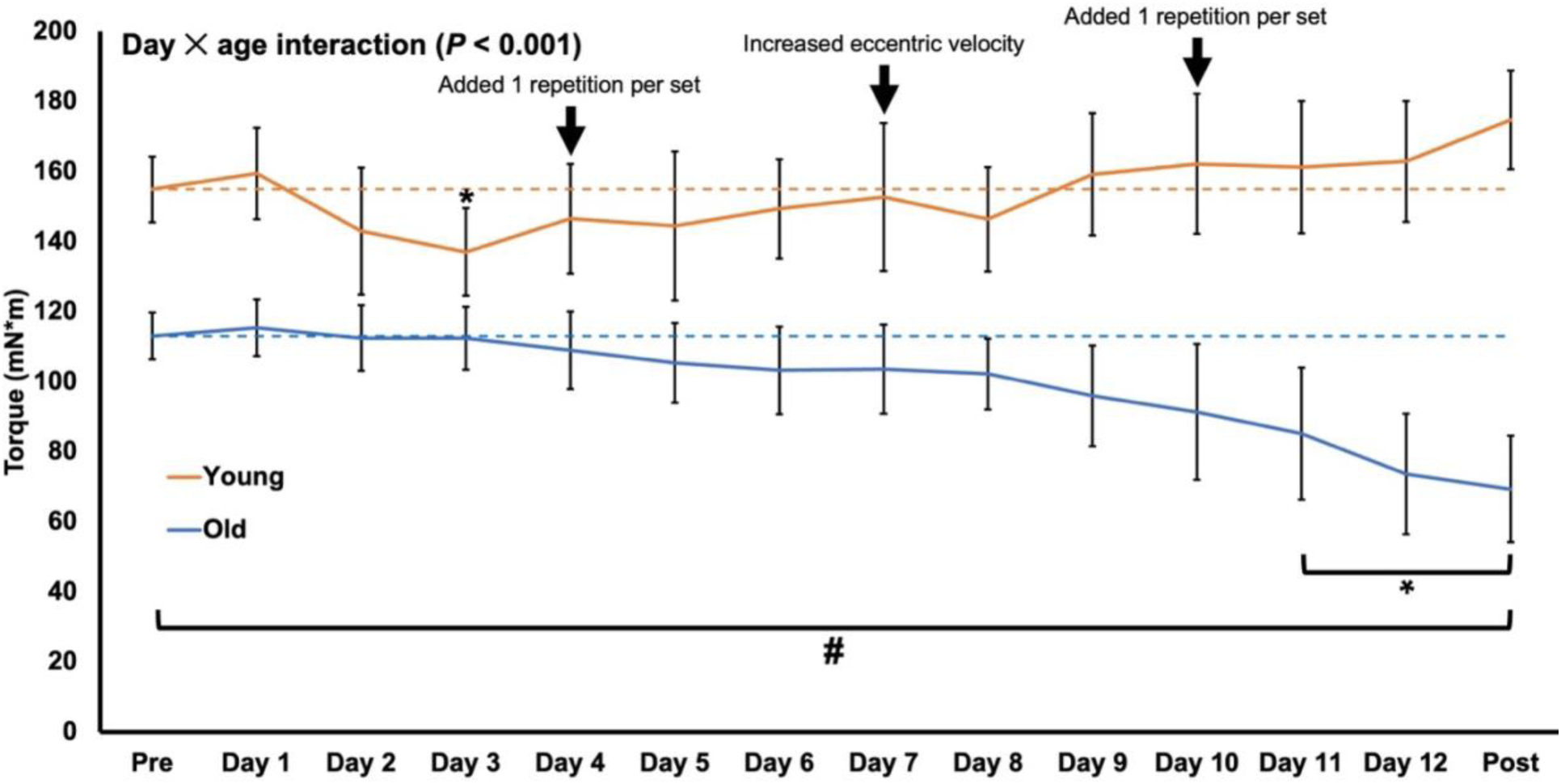
100 Hz isometric active torque at a 90° ankle angle (collected prior to training that day) throughout the eccentric training period for young (n = 10) and old rats (n = 11). Training occurred 3 days per week (Monday, Wednesday, Friday) for 4 weeks. The dashed horizontal lines represent torque at Pre. *Significant difference from Pre (*P* < 0.05). #Significant difference between young and old (*P* < 0.05). Data are presented as mean ± standard deviation.

### Changes in the passive torque-angle relationship between young and old and pre to post-training

Full ANOVA results for the passive torque-angle relationship are displayed in Supplemental Figure S7. There was a training × age × angle interaction for passive torque. For all ages and time points, passive torque increased as the ankle angle became more acute (*P* < 0.001-0.043) (Figure 10). Pre-training, old rats had 62-191% greater resting passive torque than young at all ankle angles (*P* < 0.001) (Figure 10). Post-training, old rats had 35-128% greater resting passive torque than young at all ankle angles (*P* < 0.001-0.005) except for 105° in which there was no difference between young and old (*P* = 0.814), and 110° in which old rats had lower passive torque than young (*P* <0.001) (Figure 10). For young rats, passive torque only increased 110% at 110° from pre- to post-training (*P* = 0.021). For old rats, passive torque decreased 172% at 110° and 28% at 105° from pre- to post-training (*P* < 0.01) and increased 24-51% at angles 90° to 70° pre to post-training (*P* < 0.001-0.017) (Figure 10).

**Figure 10:**
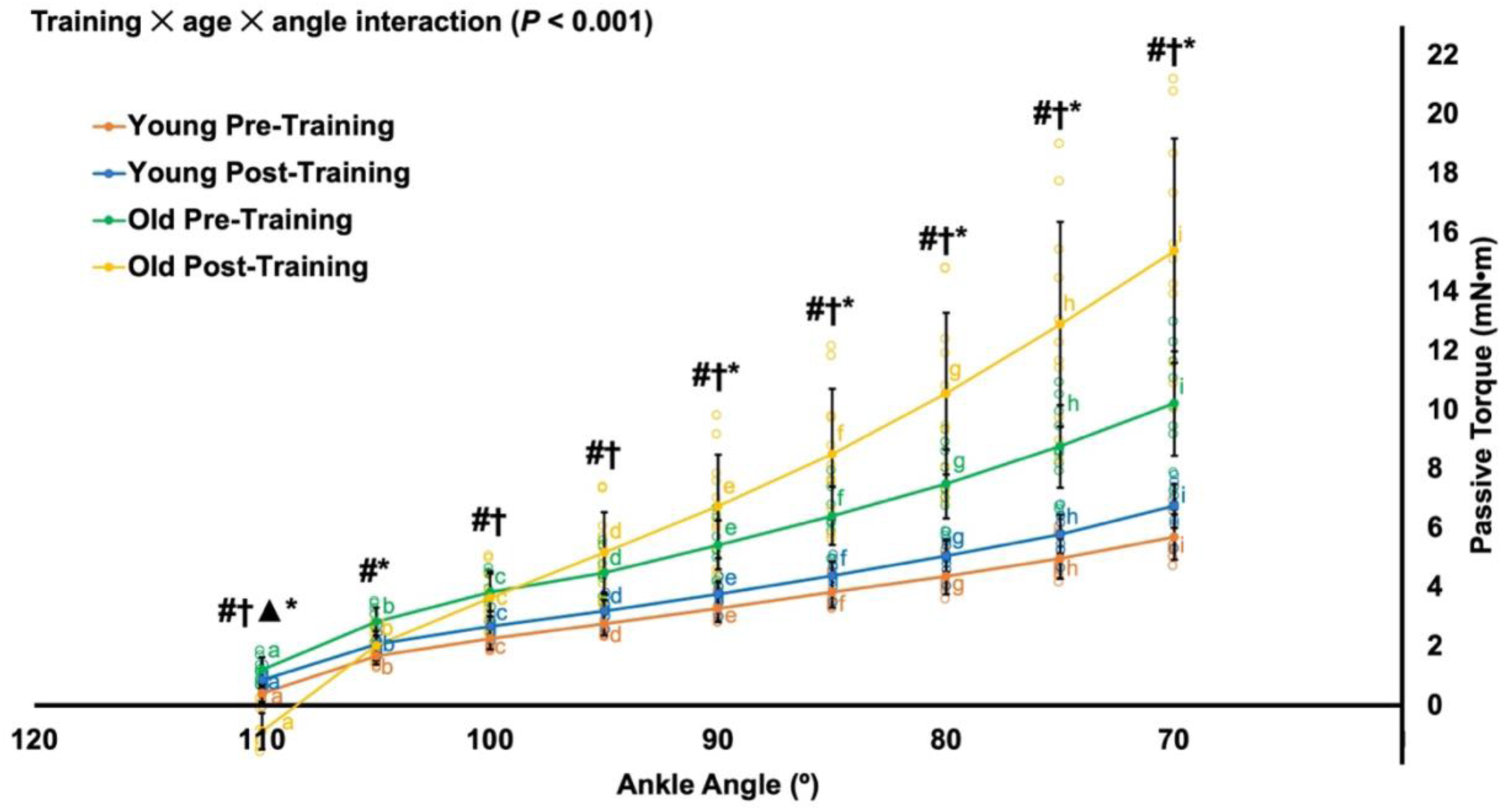
Passive torque-angle relationship of the plantar flexors pre- and post-training in young (n = 10) and old rats (n = 11). #Significant difference between young and old at pre-training (*P* < 0.05). †Significant difference between young and old at post-training (*P* < 0.05). ▲Significant difference between young pre- and post-training (*P* < 0.05). *Significant difference between old pre- and post-training (*P* < 0.05). Same letters indicate no significant difference between points within each color (*P* > 0.05). Data are presented as mean ± standard deviation.

### Changes in torque-angular velocity-power properties between young and old and pre to post-training

Full ANOVA results for angular velocity across the torque-angular velocity relationship are displayed in Supplemental Figure S8. There was a training × age × load interaction for angular velocity (F(2.121,40.294) = 19.686, *P* < 0.001, η_p_^2^ = 0.509). Both pre (–28 to –39%) and post-training (–61 to –78%), old rats produced slower velocities than young at each load (*P* <0.001-0.012) (Figure 11A-B). For young rats, velocity did not change at any loads from pre- to post-training (*P* = 0.325-0.990) (Figure 11A-B). For old rats, velocity decreased 43-61% at all loads from pre- to post-training (*P* <0.001-0.004) (Figure 11A-B).

**Figure 11:**
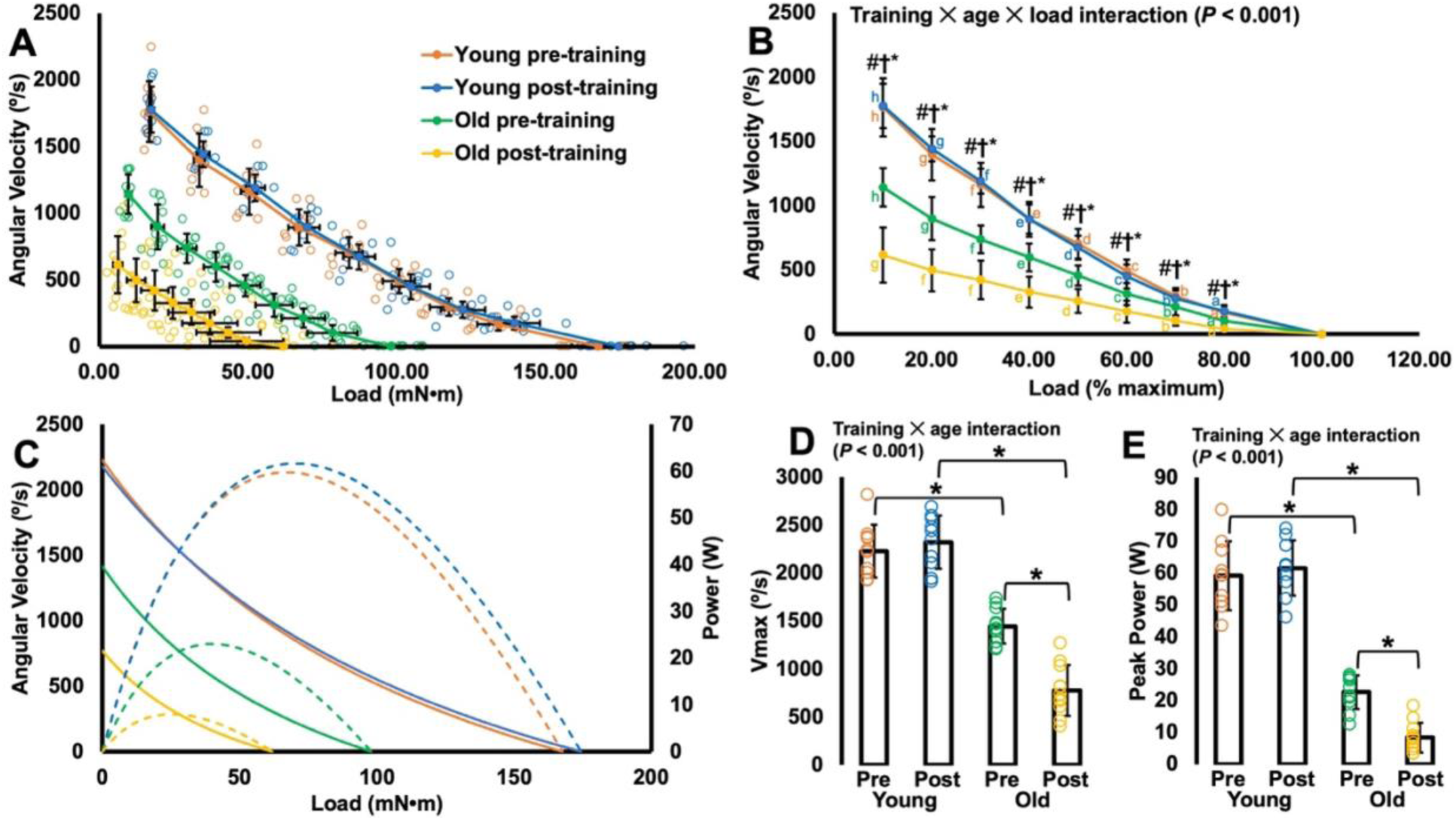
Torque-angular velocity-power properties of the plantar flexors pre- and post-training in young (n = 10) and old rats (n = 11). **A:** Angular velocity as a function of absolute load with individual data points. **B:** Angular velocity as a function of relative load. †Significant difference in angular velocity between young and old at post-training (*P* < 0.05). *Significant difference between old pre- and post-training (*P* < 0.05). Same letters indicate no significant difference between points within each color (*P* > 0.05). **C:** Fitted torque-angular velocity (solid lines; left axis) curves using the Hill equation with peak power (dashed lines; right axis) computed by multiplying angular velocity by load. **D:** Maximum shortening velocity (V_max_) extrapolated from the fitted curves in C. *Significant difference between indicated points (*P* < 0.05). **E:** Peak power obtained from the fitted curves in C. *Significant difference between indicated points (*P* < 0.05). Data are presented as mean ± standard deviation.

Full ANOVA results for V_max_, peak power, torque and velocity at peak power, curvature of the torque-velocity relationship, and the *a* and *b* coefficients of the Hill equation are shown in Supplemental Figure S5.

For V_max_, there was a training × age interaction (F(1,19) = 33.549, *P* < 0.001, η_p_^2^ = 0.638). V_max_ was slower in old than young rats both pre (–35%) and post-training (–67%) (*P* < 0.001) (Figure 11C-D). In young rats, V_max_ did not change from pre- to post-training (*P* = 0.343). In old rats, V_max_ decreased 46% from pre- to post-training (*P* < 0.001) (Figure 11C-D).

For peak power, there was a training × age interaction (F(1,19) = 21.402, *P* < 0.001, η_p_^2^ = 0.530). Old rats were less powerful than young rats both pre (–62%) and post-training (–86%) (*P* < 0.001) (Figure 11C and E). In young rats, peak power did not change from pre- to post-training (*P* = 0.372) (Figure 11C and E). Old rats, however, became 63% less powerful from pre- to post-training (*P* < 0.001) (Figure 11C and E). This age-related decrease in peak power seemed to be due to decreases in both torque (F(1,19) = 28.156, *P* < 0.001, η_p_^2^ = 0.597) and velocity (F(1,19) = 29.021, *P* < 0.001, η_p_^2^ = 0.604) at peak power, which both exhibited a training × age interaction. Both torque (–42 to –63%) and velocity (–35 to –65%) at peak power were less in old than young rats pre- and post-training (*P* < 0.001). Torque and velocity at peak power also did not differ from pre- to post-training for young rats (*P* = 0.504-0.581), but decreased 36% and 46%, respectively, from pre- to post-training for old rats (*P* < 0.001).

There were no differences in curvature of the torque-velocity relationship between young and old rats (F(1,19) = 0.089, *P* = 0.769) nor between pre- and post-training (F(1,19) = 0.523, *P* = 0.478). For the *a* and *b* coefficients, there were effects of age (*a*: F(1,19) = 11.560, *P* = 0.003, η_p_^2^ = 0.378; *b*: F(1,19) = 14.621, *P* = 0.001, η_p_^2^ = 0.435) but not training (*a*: F(1,19) = 0.664, *P* = 0.425; *b*: F(1,19) = 3.127, *P* = 0.093), such that the *a* and *b* coefficients were 48% and 45% less, respectively, in old than young rats.

## Discussion

The purpose of this study was to investigate age-related differences in serial sarcomerogenesis and adaptations in mechanical function of the plantar flexors following 4 weeks of maximal isokinetic eccentric training. The main finding was that sarcomerogenesis in old rats was impaired, leading to unfavourable mechanical adaptations, while young rats showed favourable architectural remodeling and mechanical adaptations. We hypothesized that SSN would increase in the plantar flexor muscles of young rats, and that this would correspond to improvements in torque production, V_max_, and peak power, and a reduction in resting passive tension. We expected old rats to experience a smaller magnitude of serial sarcomerogenesis, alongside smaller improvements in mechanical function as compared with young. Our hypotheses were partly correct. In the soleus, LG, and MG, old rats had a lower SSN than young. Pre-training, old rats were also weaker, slower, less powerful, and had greater passive tension than young rats. Following training, young rats exhibited serial sarcomerogenesis, while old rats exhibited clear deficits in the ability to add sarcomeres in series, most noticeably in the soleus. Young rats experienced small improvements in mechanical function, including improved torque production and an apparent widening of the torque-angle relationship’s plateau region, but with few alterations to the passive torque-angle and torque-angular-velocity-power relationships. Old rats, conversely, became weaker, slower, and less powerful, and exhibited a steeper rise in passive tension with increasing muscle stretch following training.

### Age-related differences in serial sarcomere number

The SSN, SL, and FL values (Figures 3-5) we observed for the rat soleus, LG, and MG are consistent with previous studies (10,18,49,58–60). In line with previous reports from Power et al. (10), we observed 14% fewer serial sarcomeres in the MG of old compared to young rats, with 14% shorter fascicles and no significant difference in SL at a 90° ankle angle. We can now expand this age-related reduction in SSN to the rat soleus and LG. In the untrained soleus, old rats had 9% fewer serial sarcomeres, with 13% shorter fascicles and 3% shorter SLs at 90°. In the untrained LG, old rats had 7% fewer serial sarcomeres, with 10% shorter fascicles and 3-5% shorter SLs in the middle and distal regions of the muscle at 90°. Studies on humans have well-documented that muscle mass/size declines progressively in old age (3,5), with ultrasonographic measurements on humans indicating reductions in both FL and fascicle pennation angle contribute to this loss (6–8). Coupled with our observation of 14-28% lower muscle wet weights in old compared to young rats (Figure 2), our findings confirm previous reports in animals (9,10) that reductions in FL contribute to the age-related loss of muscle mass due to the loss of serial sarcomeres.

### Age-related differences in mechanical function pre-training

The isometric torque values we observed in the young rats are within those reported previously for the rat plantar flexors *in vivo* for 3-5-month-old rats (52,61). Pre-training, young rats produced greater isometric tetanic torque at 70° (longer muscle lengths) than 90°, while old rats produced greater torque at 90° (shorter muscle lengths) than 70° (Figure 6). Old rats producing more optimal torque at a shorter muscle length aligns with them having a lower SSN than the young rats, as moving to 70° would stretch their sarcomeres onto the descending limb of the force-length relationship, where fewer force-producing cross-bridges would form (19,21,46). Conversely, young rats, with a greater SSN, experienced more optimal cross-bridge formation at a longer muscle length. Importantly, our SL measurements were recorded with the knee fully extended and the ankle at 90°, rather than at the muscles’ optimal lengths for force production.

Therefore, the shorter SLs we measured in old compared to young do not reflect a shorter optimal SL per se, and since vertebrates exhibit little variability in optimal SL (62), no age-related change in optimal SL was expected. With that said, if old rats produced optimal active torque at 90° while young rats produced optimal active torque at 70°, the shorter SLs recorded for the soleus and LG in old rats may in fact allude to a shorter optimal SL. In support of this theory, Power (10) observed an average (though not significant) 4% shorter optimal SL of the MG in old than young rats. It is also important to note that the 3% decreases in SL we observed would amount to about 66 nm. With a laser wavelength of 635 nm, the resolution would be about 300 nm, making detecting a 66 nm change in SL have some possibility for error. Regardless, our observations warrant further research on potential age-related alterations to optimal SL.

Prior to training, old rats produced 27% lower torque at 90° and 42% lower torque at 70° than young rats during maximal activation, which is consistent with a previous comparison between 6-month-old rats and 30-month-old rats for the dorsiflexors (63) along with a wealth of studies comparing maximum torque between young and older humans for various muscle groups (for reviews see: 3,5,13,62). The greater age-related torque deficit at 70° than 90° can also likely be explained by the lower SSN in old rats, with only young rats producing optimal torque at 70°. When normalizing maximum torque to the combined wet weight of the gastrocnemii and soleus, the difference in torque between young and old rats pre-training at 90° was eliminated and at 70° was reduced by 19%, indicating lower muscle mass contributed to a large portion of the age-related strength loss we observed. We also observed lower torque in old compared to young rats at submaximal stimulation frequencies of 20 to 80 Hz. These reductions in torque (both maximal and submaximal) are a result of a combined loss of motor units and muscle fibre cross-sectional area with age, which together manifest as a loss of muscle mass (5). There is, in particular, a preferential loss of type II fibres in old age (65) or at least a preferential loss of type II fibre area such that these fibres become weaker (5). We also observed a lower F_50_ in old rats, with 50% of maximum torque being developed at ∼30 Hz in old but ∼40 Hz in young. The lower F_50_ in old rats can be explained by the 25% longer half-relaxation time we observed in old than young rats, as slower relaxation allows twitches to summate at a lower stimulation frequency. These age-related differences in F_50_ and twitch half-relaxation time likely reflect a greater proportion of type I fibres in old rats, consistent with a previous study showing a greater proportion of type I fibres in the MG of old (∼28 months) compared to middle-aged (∼13 months) rats (65).

Aligning with previous reports in humans (5), old rats produced slower angular velocities at all relative isotonic loads, a slower V_max_, and a lower peak power than young rats (Figure 11). The slower velocities may partly reflect a shift toward a greater whole muscle type I (slow):type II (fast) fibre area (5), but may also be partly due to the lower SSN. When force-producing structures are arranged in series, their forces are shared while their individual excursions are additive. As velocity equals excursion over time, SSN (and optimal FL) is proportional to a muscle’s V_max_ (13,66,67). Accordingly, Thom et al. (12) observed that MG FL measured via ultrasound accounted for almost half the age-related difference in V_max_ in humans. The losses of torque and velocity together contributed to the 62% lower peak power we observed in the old rats compared to young, as power is the product of both torque and velocity. It appears the loss of torque may have contributed slightly more than velocity to age-related power loss, as torque at peak power was 42% less while velocity at peak power was 35% less in old than young rats.

Lastly, old rats incurred greater plantar flexor passive torque than young rats across all joint angles tested (70° to 110°). While it may seem reasonable to speculate that the fewer serial sarcomeres in old rats caused sarcomeres to be more overstretched, and thus produce greater passive forces, this may not be the case as SL was the same or shorter in old rats than young rats as measured at 90°. Similarly, Power et al. (10) observed that a lower SSN in the old rat MG could not explain elevated MG passive tension in old rats, instead attributing elevated passive tension to a remodelling of collagen slack length. Therefore, the greater passive torque in old compared to young rats is likely a result of age-related differences in the ECM. Age-related increases in ECM area and collagen content have been observed in mice (68–71) and humans (41). Muscles of old mice also have more collagen crosslinks than young mice (69,72–75), in particular with more advanced glycation end products (69,74,76), which increases the stiffness and strength of collagen fibrils, causing passive tension to be elevated when stretched (77). It follows that collagen crosslinking is largely responsible for elevated passive stiffness in aged muscle (78–81).

### Changes in mechanical function following eccentric training in young rats

The maximum active plantar flexion torque of young rats at 90° increased 13% from pre- to post-training. This increase is less than the 25% increase in maximum plantar flexion torque at 90° following eccentric training observed by Ochi et al. (52). Their results may differ from ours because they used 3-month-old rats while we used 9-month-old rats, and Rader et al. (63) showed 3-month-old rats experience greater training-induced increases in strength than 6-month-old rats, likely due to the concomitant effect of maturation-related growth on strength. Notably, only maximum torque at 90° increased in the present study while torque at 70° remained unchanged. This joint angle-dependent increase in strength likely reflects broadening of the torque-angle relationship’s plateau region, which is expected to occur alongside serial sarcomerogenesis. With more sarcomeres in series, individual sarcomeres can stay closer to optimal SL at more joint angles throughout the range of motion (14,16,82). The increase in torque at 90° pre- to post-training appeared to be largely due to an increase in muscle mass, as normalizing torque to total muscle wet weight of the soleus and gastrocnemii eliminated the difference from pre- to post-training. The increases in soleus and MG muscle mass could reflect the observed increases in SSN, but could also be partly from increases in fibre cross-sectional area (i.e., parallel sarcomere number), which has been observed with eccentric training in rats (52) and humans (83). While an increase in SSN may also be expected to increase V_max_ and peak power (see previous section) (16,84,85), those remained unchanged following eccentric training in our young rats. Therefore, while serial sarcomerogenesis altered static contractile function in the present study, it did not appear to impact dynamic contractile function.

A decrease in passive torque can also be expected alongside serial sarcomerogenesis due to less stretch of individual sarcomeres and thereby less passive tension development at a given joint angle, assuming joint moment arms remain the same (16). However, we observed the opposite, with a slight upward shift in the passive torque-angle relationship for young rats (Figure 10). Increases in tendon passive stiffness (30,86) and ECM remodelling (53,87–89) are both common following eccentric exercise, and both could contribute to greater passive tension of a muscle-tendon unit. Therefore, adaptations in these non-contractile structures may have concealed the effect of a greater SSN on the passive torque-angle relationship.

### Changes in mechanical function following eccentric training in old rats

In stark contrast to what we observed in young rats, old rats exhibited reductions in active plantar flexion torque at 90° and 70° and at each stimulation frequency, a slowing of angular velocity at each relative load, slowing of V_max_, and a reduction in peak power. An upward shift in the passive torque-angle relationship also alluded to greater stiffening of the ECM following eccentric training compared to young rats. Rader et al. (63) also observed an inability of old rats to adapt to training, with old rats exhibiting no change in maximum isometric force following 4 weeks of dorsiflexor stretch-shortening cycle training. The reduction in torque in the present study could not entirely be accounted for by a loss of muscle mass, as muscle wet weights of the trained leg were not less than the untrained leg for old rats, and a reduction in torque was still observed when normalizing to total muscle wet weight (Figure 7). These reductions in torque were also unlikely to be caused by an additional 4 weeks of natural aging, as two 34-month-old untrained rats (i.e., 1 month older than the trained rats post-training) produced torque values within those recorded for the trained cohort pre-training (Supplemental Figure S9). Therefore, the impairments in mechanical function that we observed appear to be due to an age-related dysfunctional adaptation to the training protocol. Reasons for this dysfunction are explored in the next section.

### Eccentric training induced serial sarcomerogenesis in young rats but revealed dysfunctional remodelling in old rats

Due to the large mechanical tension borne by the muscle during active stretch, training programs emphasizing eccentric contractions are often used to stimulate serial sarcomerogenesis (18–21,49). Serial sarcomerogenesis is believed to occur to better optimize actin-myosin overlap and reduce passive tension in a stretched position (16,46,56). As well, eccentric exercise can stretch sarcomeres to the point that Z-lines are damaged, and adapting by adding sarcomeres in series allows a muscle to stretch further without this sarcomere damage occurring on subsequent bouts (90–93).

The magnitude of SSN increase we observed in the young rat soleus and MG is similar to that observed previously in the soleus of 18-week-old rats following weighted downhill running training (18), and in the vastus intermedius of 5-6-month-old rats following bodyweight downhill running training (19,20). In the present study, the SSN of young and old rats exhibited different responses to eccentric training (Figures 3-5). In young rats, eccentric training resulted in a 4% increase in soleus SSN from the untrained to the trained leg, while old rats saw no difference in soleus SSN. For the MG, though there was no significant training × age interaction, and an effect of training showed an overall 4% increase from the untrained to trained leg, it should be noted that with the age groups separated, young rats showed a significant 6% increase in SSN (P = 0.01; two-tailed, paired t-test) while old rats showed no significant difference in SSN (P = 0.39) and on average only a 2% increase. Also, the LG of young rats did not differ in SSN between the untrained and trained leg, while the LG of old rats experienced a 7% decrease in SSN. Altogether, young rats showed the expected response of adding serial sarcomeres with eccentric training while old rats did not exhibit this adaptation, even experiencing some sarcomere loss.

Old rats may have experienced blunted serial sarcomerogenesis due to their muscles being more susceptible to eccentric exercise-induced damage than young, and thus requiring longer periods to recover. A hallmark of eccentric exercise-induced damage is a long-lasting deficit in maximum torque production. McBride et al. (36) exposed the ankle dorsiflexors of old (32 months) and young (6 months) rats to four sets of six eccentric contractions. While the initial drop in maximum torque (∼30%) they observed was similar between ages, young rats recovered by 5 days post-exercise while old rats remained deficient in torque by 18% at 5 days post-exercise (36). Furthermore, following a subsequent bout performed two weeks later, young rats exhibited no deficit in maximum torque, while old rats exhibited a similar drop in torque to the first bout. Therefore, there is an impaired ability to both recover and adapt following one bout of eccentric exercise in old rats. Similar findings have also been observed in mice (35,94–96) and humans (37–39). Considering Figure 9 (changes in 100 Hz torque throughout the training period), this inability for old rats to recover from and adapt to eccentric exercise appeared to occur in the present study too. Our young rats exhibited an initial drop in maximum torque during the first week (day 1-3) of training, then subsequently recovered and even gradually increased beyond pre-training maximum torque, even when the training stimulus was enhanced at days 4, 7 and 10. Old rats, conversely, had a gradual drop in maximum torque throughout the entire training period, which was accelerated by adding faster eccentric contractions (which are typically more damage-inducing (44,45)) starting at day 7. A recent study comparing old and young mice observed similar findings, with young mice becoming 20-36% stronger but old mice becoming 18% weaker following 5 weeks of high-intensity eccentric training 1 day/week (97). One explanation in the present study may be the lower pre-training SSN in old rats compared to young. Assuming similar series elastic compliance, with fewer serial sarcomeres, old rats’ individual sarcomeres would have undergone a greater stretch amplitude for the same joint rotation, thereby incurring more damage and requiring a longer recovery period (36,44,98,99). Old rats incurring greater damage may be supported by the greater SL SD in the trained compared to untrained MG of our old rats (Supplemental Figure S10F), as greater SL non-uniformity may correspond to greater ultrastructural damage following long-term eccentric training (100).

For serial sarcomerogenesis to occur, the mechanical stretch must be transmitted from the ECM to transmembrane complexes and other cytoskeletal proteins of the muscle fibre (8,101–104). Deformation of those proteins activates biochemical events that regulate sarcomeric gene transcription and protein synthesis for the addition of new sarcomeres; these processes collectively are known as mechanotransduction (8,105). In our recent review, we identified age-related changes in mechanotransduction pathways that may limit serial sarcomerogenesis, including impaired mechanistic target of rapamycin and insulin-like growth factor 1 signaling, increased myostatin activity, reduced activation of serum response factor, increased protein degradation by muscle ring finger protein, and reduced satellite cell content and function (see Figure 3 of Hinks et al. (8)). These age-related changes may be partly responsible for the poor adaptability observed following eccentric training in old rats. Additionally, as discussed earlier, the ECM tends to accumulate more collagen and collagen crosslinks with age, giving aged muscle a more fibrotic appearance and increased passive stiffness (77). Pre-training collagen content and packing density are negatively associated with hypertrophic outcomes (42,43), likely because a stiffer, more crosslinked ECM limits the mechanical stimuli imposed on the muscle contractile tissue. To that point, Rahman et al. (102) observed that impaired ECM remodelling limited the regenerative response following damage to the tibialis anterior induced by cardiotoxin in old compared to young mice. Furthermore, Baumann et al. (97) compared gene profiles of young and old mice following 5 weeks of eccentric training, and found that while young and old mice both exhibited upregulation of genes associated with sarcomere organization and assembly, only old mice exhibited upregulation of genes associated with ECM remodelling, likely indicative of fibrosis. This remodeling of the local connective tissue environment in old may blunt the ability for a mechanical signal to be transduced into a biochemical response (8,107), limiting serial sarcomerogenesis. In-depth investigation of ECM content and architecture, and mechanotransduction protein content and gene expression are beyond the scope of the present study, thus further research is needed to explore these potential causes of blunted sarcomerogenesis in more detail.

If we had used slower eccentric contractions, a submaximal eccentric torque, or more recovery days between training sessions, it is possible we would have mitigated the development of damage and provided better conditions for adaptation in the old rats (108). However, considering again the findings of McBride et al. (36), who employed a less intense eccentric exercise protocol (80% of maximum force, four sets of only six repetitions with 5 minutes between sets) and still observed not even half recovery of maximum torque 5 days post-exercise, using a less intense eccentric training protocol may not have altered the outcome of our study. With that said, studies on older men and women (average age 65-74 years) have observed increases in FL as measured by ultrasound, strength, V_max_, and power following submaximal (working at ∼50-75% of 1-repetition maximum) eccentric training 2-3 days per week for 7-16 weeks (30–32,109). Therefore, this age-related inability to recover from and adapt to eccentric training may be more pronounced in rodents than humans. One notable difference is that studies on rodents use controlled, electrically stimulated activations while studies on humans use voluntary contractions. Though speculative, a potential explanation may be that the neural component of maximal voluntary muscle activation in humans could offset the impact on the muscle itself. Future studies should explore modifications to eccentric training programs to better elucidate the optimal dose/stimulus for serial sarcomerogenesis in old age (108).

### Conclusions

We found that young and old rats exhibited opposing adaptations to eccentric training. In young rats, eccentric training increased SSN, improved torque production, and minimally altered the passive torque-angle and torque-velocity-power relationships. Old rats, conversely, exhibited blunted sarcomerogenesis, reductions in torque, velocity, and power production, and an upward shift in the passive torque-angle relationship following training. Old rats may have been more susceptible to muscle damage induced by the eccentric exercise, preventing them from recovering and adapting between training sessions, leading to dysfunctional remodeling. Future studies should explore potential causes for blunted sarcomerogenesis following eccentric exercise in old age, and modifications to eccentric training to better understand the optimal dose/stimulus for beneficial adaptations in old age.

## List of abbreviations

ANOVA: analysis of variance
ECM: extracellular matrix
F_50_: frequency at which 50% of maximum torque is produced
FL: fascicle length
HRT: half-relaxation time
LG: lateral gastrocnemius
MG: medial gastrocnemius
SL SD: sarcomere length standard deviation
SL: sarcomere length
SSN: serial sarcomere number
V_max_: maximum shortening velocity

## Acknowledgements

This project was supported by the Natural Sciences and Engineering Research Council of Canada (NSERC). No conflicts of interest, financial or otherwise, are declared by the authors. We would like to thank Dr. Walter Herzog for providing comments on a previous version of this manuscript.

## Ethics statement

Approval was given by the University of Guelph’s Animal Care Committee and all protocols followed CCAC guidelines (AUP #4905).

## Data availability

All data generated or analysed during the study are available from the corresponding author upon request.

## Author contributions

A.H. and G.A.P. conceived and designed research; A.H., M.A.P., and B.S.N. carried out animal husbandry and training; A.H. performed experiments; A.H. analyzed data; A.H., M.A.P, B.S.N., and G.A.P. interpreted results of experiments; A.H. prepared figures; A.H. and G.A.P. drafted manuscript; A.H., M.A.P., B.S.N., and G.A.P. edited and revised manuscript; A.H., M.A.P., B.S.N., and G.A.P. approved final version of manuscript.

---

**Supplemental Figure S9:**
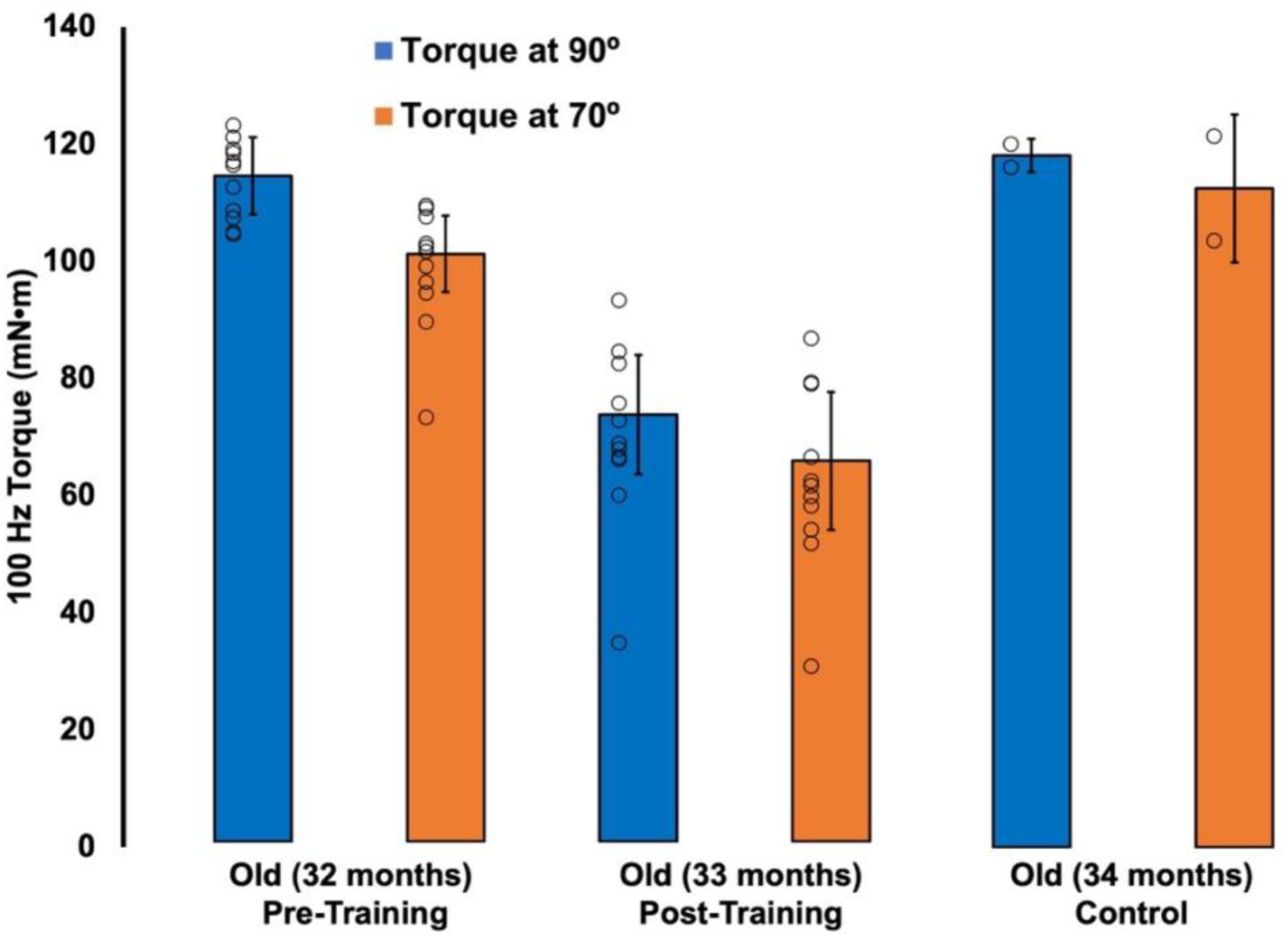
Maximum plantar flexion torque values for old rats at pre (32 months old) and post-eccentric training (33 months old) (n = 11), and untrained rats tested one month later (34 months old) (n = 2).

**Supplemental Figure S10:**
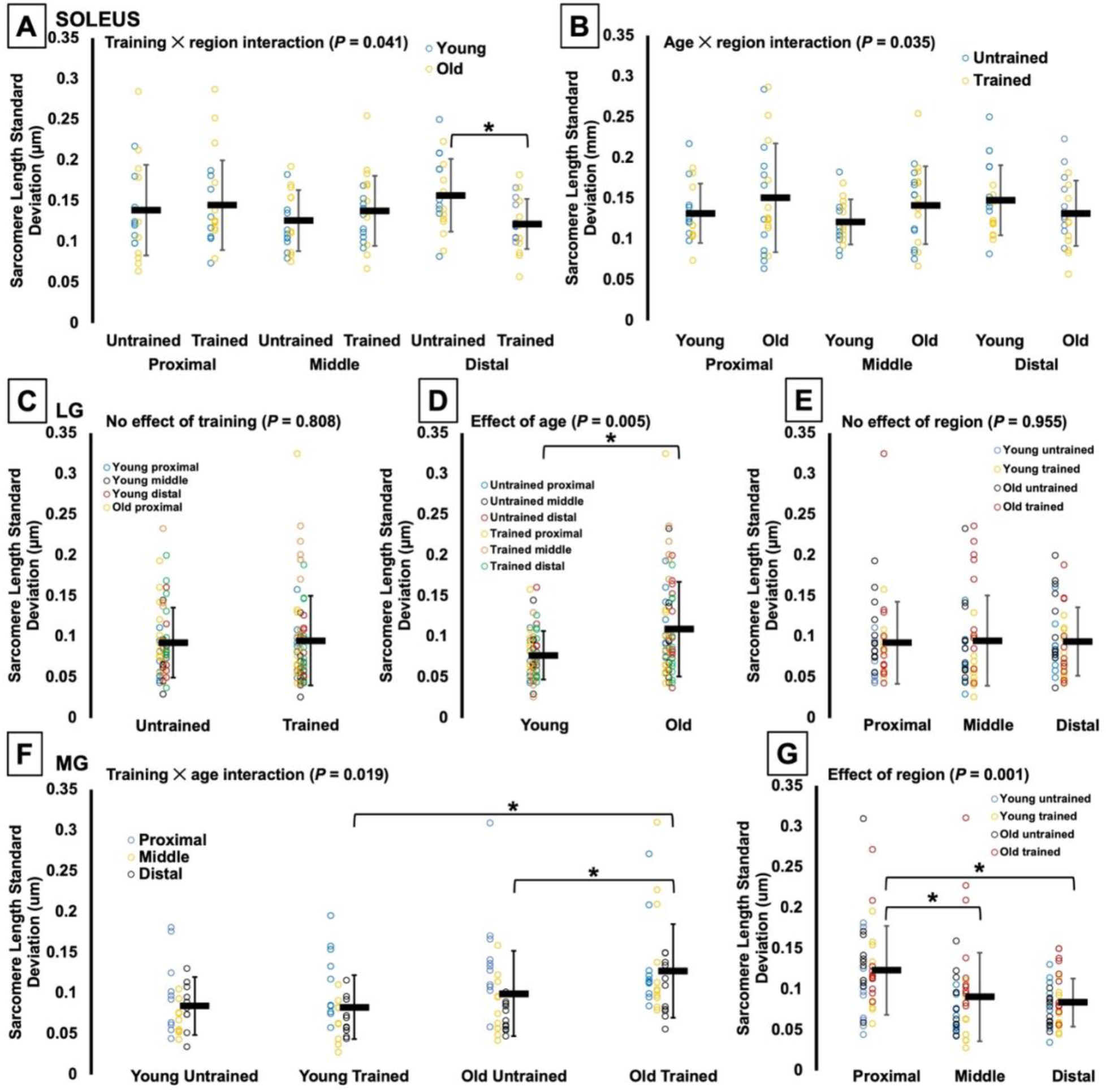
Sarcomere length standard deviation of the soleus (**A and B**), lateral gastrocnemius (LG) (**C-E**), and medial gastrocnemius (MG) (**F and G**) in the trained and untrained legs of young and old rats across the proximal, middle, and distal muscle regions. *Significant difference between indicated points (P < 0.05).

