## Supplementary material for "Age-related blunting of serial sarcomerogenesis and mechanical adaptations following 4 weeks of maximal eccentric resistance training": Figures S1-8

**Supplemental Figure S1: Two-way ANOVA (Training × Age) results for Muscle Wet Weight**

| Effect | Muscle | F | P | $\eta_p^2$ |
|---|---|---|---|---|
| **Training** | Soleus | 7.121 | **0.015*** | 0.273 |
| | LG | 1.367 | 0.257 | 0.067 |
| | MG | 16.677 | **<0.001*** | 0.467 |
| **Age** | Soleus | 21.476 | **<0.001*** | 0.531 |
| | LG | 51.681 | **<0.001*** | 0.731 |
| | MG | 109.960 | **<0.001*** | 0.853 |
| **Training × Age** | Soleus | 1.839 | 0.191 | 0.088 |
| | LG | 0.964 | 0.339 | 0.048 |
| | MG | 0.205 | 0.656 | 0.011 |

LG = lateral gastrocnemius; MG = medial gastrocnemius; *Significant effect or interaction

**Supplemental Figure S2: Three-way ANOVA (Training × Age × region) results for serial sarcomere number, fascicle length, and sarcomere length**

| | | Training | | | Region | | | Age | | |
|---|---|---|---|---|---|---|---|---|---|---|
| | | F | P | $\eta_p^2$ | F | P | $\eta_p^2$ | F | P | $\eta_p^2$ |
| **Soleus** | **SSN** | 2.163 | 0.158 | 0.102 | 0.531 | 0.592 | 0.027 | 138.404 | **<0.001*** | 0.879 |
| | **FL** | 0.103 | 0.752 | 0.005 | 2.090 | 0.138 | 0.099 | 120.350 | **<0.001*** | 0.864 |
| | **SL** | 1.686 | 0.210 | 0.082 | 4.486 | **0.019*** | 0.191 | 2.709 | 0.116 | 0.125 |
| **LG** | **SSN** | 0.048 | 0.829 | 0.003 | 36.693 | **<0.001*** | 0.659 | 30.568 | **<0.001*** | 0.617 |
| | **FL** | 0.830 | 0.374 | 0.042 | 11.495 | **<0.001*** | 0.377 | 56.347 | **<0.001*** | 0.748 |
| | **SL** | 1.637 | 0.216 | 0.079 | 67.058 | **<0.001*** | 0.779 | 10.285 | **0.005*** | 0.351 |
| **MG** | **SSN** | 7.823 | **0.011*** | 0.292 | 31.074 | **<0.001*** | 0.621 | 36.312 | **<0.001*** | 0.656 |
| | **FL** | 10.107 | **0.005*** | 0.347 | 16.340 | **<0.001*** | 0.462 | 51.640 | **<0.001*** | 0.731 |
| | **SL** | 1.395 | 0.252 | 0.068 | 49.011 | <**0.001*** | 0.721 | 0.002 | 0.961 | 0.000 |

| | | Training × Age | | | Region × Age | | | Training × Region | | | Training × Region × Age | | |
|---|---|---|---|---|---|---|---|---|---|---|---|---|---|
| | | F | P | $\eta_p^2$ | F | P | $\eta_p^2$ | F | P | $\eta_p^2$ | F | P | $\eta_p^2$ |
| **Soleus** | **SSN** | 10.099 | **0.005*** | 0.347 | 0.042 | 0.959 | 0.002 | 0.809 | 0.453 | 0.041 | 0.038 | 0.963 | 0.002 |
| | **FL** | 1.483 | 0.238 | 0.072 | 0.072 | 0.930 | 0.004 | 0.330 | 0.721 | 0.017 | 0.505 | 0.608 | 0.026 |
| | **SL** | 4.896 | **0.039*** | 0.205 | 1.683 | 0.201 | 0.081 | 1.011 | 0.361 | 0.051 | 1.165 | 0.316 | 0.058 |
| **LG** | **SSN** | 7.831 | **0.011*** | 0.292 | 1.966 | 0.154 | 0.094 | 0.026 | 0.975 | 0.001 | 0.281 | 0.757 | 0.015 |
| | **FL** | 10.238 | **0.005*** | 0.350 | 1.685 | 0.199 | 0.081 | 0.050 | 0.951 | 0.003 | 0.193 | 0.825 | 0.010 |
| | **SL** | 0.000 | 0.986 | 0.000 | 5.805 | **0.009*** | 0.234 | 1.240 | 0.299 | 0.061 | 1.423 | 0.254 | 0.070 |
| **MG** | **SSN** | 2.244 | 0.151 | 0.106 | 0.807 | 0.454 | 0.041 | 1.096 | 0.344 | 0.055 | 2.017 | 0.147 | 0.096 |
| | **FL** | 0.772 | 0.391 | 0.039 | 1.063 | 0.339 | 0.053 | 0.828 | 0.436 | 0.042 | 2.099 | 0.141 | 0.099 |
| | **SL** | 1.621 | 0.218 | 0.079 | 2.928 | 0.066 | 0.134 | 5.267 | **0.012*** | 0.217 | 0.050 | 0.939 | 0.003 |

SSN = serial sarcomere number; FL = fascicle length; SL = sarcomere length; LG = lateral gastrocnemius; MG = medial gastrocnemius; *Significant effect or interaction

**Supplemental Figure S3: Three-way ANOVA (Training × Age × Angle) for absolute and mass-specific 100 Hz torque at 90° and 70°**

| **ABSOLUTE 100 HZ TORQUE (mN•m)** | | | |
|---|---|---|---|
| **Effect** | **F** | **P** | $\eta_p^2$ |
| **Training** | 25.878 | **<0.001*** | 0.577 |
| **Angle** | 6.149 | **0.023*** | 0.245 |
| **Age** | 390.038 | **<0.001*** | 0.954 |
| **Training × Age** | 103.541 | **<0.001*** | 0.845 |

| | | | |
|---|---|---|---|
| **Age × Angle** | 69.106 | **<0.001*** | 0.784 |
| **Training × Angle** | 1.477 | 0.239 | 0.072 |
| **Training × Age × Angle** | 21.785 | **<0.001*** | 0.534 |
| **MASS-SPECIFIC 100 HZ TORQUE (mN•m/g)** | | | |
| **Effect** | **F** | **P** | **$\eta_p^2$** |
| **Training** | 34.143 | **<0.001*** | 0.642 |
| **Angle** | 74.559 | **0.002*** | 0.411 |
| **Age** | 151.078 | **<0.001*** | 0.888 |
| **Training × Age** | 46.255 | **<0.001*** | 0.709 |
| **Age × Angle** | 75.401 | **<0.001*** | 0.799 |
| **Training × Angle** | 0.178 | 0.678 | 0.009 |
| **Training × Age × Angle** | 19.868 | **<0.001*** | 0.511 |

*Significant effect or interaction

**Supplemental Figure S4: Three-way ANOVA (Age × Training × Frequency) for the torque-frequency relationship**

| **Effect** | **F** | **P** | **$\eta_p^2$** |
|---|---|---|---|
| **Training** | 37.564 | **<0.001*** | 0.664 |
| **Frequency** | 2355.617 | **<0.001*** | 0.992 |
| **Age** | 248.502 | **<0.001*** | 0.929 |
| **Training × Age** | 94.789 | **<0.001*** | 0.834 |
| **Age × Frequency** | 380.095 | **<0.001*** | 0.952 |
| **Training × Frequency** | 22.226 | **<0.001*** | 0.539 |
| **Training × Age × Frequency** | 51.410 | **<0.001*** | 0.730 |

*Significant effect or interaction

**Supplemental Figure S5: Two-way ANOVA (Training × Age) for mass-specific torque at 90°, maximum shortening velocity, peak power, torque and velocity at peak power, curvature of the torque-velocity relationship, and the *a* and *b* coefficients of the torque-velocity relationship**

| | **Training** | | | **Age** | | | **Training × Age** | | |
|---|---|---|---|---|---|---|---|---|---|
| | **F** | **P** | **$\eta_p^2$** | **F** | **P** | **$\eta_p^2$** | **F** | **P** | **$\eta_p^2$** |
| **$F_{50}$** | 2.266 | 0.149 | 0.107 | 205.562 | **<0.001*** | 0.915 | 12.021 | **0.003*** | 0.388 |

| | | | | | | | | | |
|---|---|---|---|---|---|---|---|---|---|
| ***n* coefficient** | 1.141 | 0.299 | 0.057 | 8.436 | **0.009*** | 0.307 | 0.588 | 0.453 | 0.030 |
| **Twitch HRT** | 1.729 | 0.204 | 0.083 | 45.170 | **<0.001*** | 0.704 | 4.499 | **0.047*** | 0.191 |
| **$V_{max}$** | 19.234 | **<0.001*** | 0.503 | 175.914 | **<0.001*** | 0.903 | 33.549 | **<0.001*** | 0.638 |
| **Peak power** | 10.908 | **0.004*** | 0.365 | 252.056 | **<0.001*** | 0.930 | 21.402 | **<0.001*** | 0.530 |
| **Torque at peak power** | 18.675 | **<0.001*** | 0.496 | 216.195 | **<0.001*** | 0.919 | 28.156 | **<0.001*** | 0.597 |
| **Velocity at peak power** | 20.925 | **<0.001*** | 0.524 | 207.117 | **<0.001*** | 0.916 | 29.021 | **<0.001*** | 0.604 |
| **Curvature** | 0.089 | 0.769 | 0.005 | 0.523 | 0.478 | 0.027 | 0.007 | 0.934 | 0.000 |
| ***a* coefficient** | 0.664 | 0.425 | 0.034 | 11.560 | **0.003*** | 0.378 | 3.095 | 0.095 | 0.140 |
| ***b* coefficient** | 3.127 | 0.093 | 0.141 | 14.621 | **0.001*** | 0.435 | 4.009 | 0.060 | 0.174 |

$F_{50}$ = frequency at which 50% of maximum torque was developed; HRT = half-relaxation time; $V_{max}$ = maximum shortening velocity;
*Significant effect or interaction

**Supplemental Figure S6: Two-way ANOVA (Day × Age) for torque at 90 degrees throughout training**

| **Effect** | **F** | **P** | **$\eta_p^2$** |
|---|---|---|---|
| **Day** | 5.078 | **<0.001*** | 0.220 |
| **Age** | 106.464 | **<0.001*** | 0.855 |
| **Day × Age** | 29.110 | **<0.001*** | 0.618 |

*Significant effect or interaction

**Supplemental Figure S7: Three-way ANOVA (Training × Age × Angle) for the passive torque-angle relationship**

| **Effect** | **F** | **P** | **$\eta_p^2$** |
|---|---|---|---|
| **Training** | 7.220 | **0.015*** | 0.275 |
| **Angle** | 781.073 | **<0.001*** | 0.976 |
| **Age** | 103.111 | **<0.001*** | 0.884 |

| | | | |
|---|---|---|---|
| **Training × Age** | 1.355 | 0.259 | 0.067 |
| **Age × Angle** | 123.042 | **<0.001*** | 0.866 |
| **Training × Angle** | 23.785 | **<0.001*** | 0.556 |
| **Training × Age × Angle** | 16.961 | **<0.001*** | 0.472 |

*Significant effect or interaction

**Supplemental Figure S8: Three-way ANOVA (Age × Training × Load) for velocity in the torque-angular velocity relationship**

| **Effect** | **F** | **P** | **$\eta_p^2$** |
|---|---|---|---|
| **Training** | 19.141 | **<0.001*** | 0.502 |
| **Load** | 1157.553 | **<0.001*** | 0.984 |
| **Age** | 164.760 | **<0.001*** | 0.897 |
| **Training × Age** | 19.447 | **<0.001*** | 0.506 |
| **Age × Load** | 135.026 | **<0.001*** | 0.877 |
| **Training × Load** | 12.582 | **<0.001*** | 0.398 |
| **Training × Age × Age** | 19.686 | **<0.001*** | 0.509 |

*Significant effect or interaction
